# Ultrasensitive and multiplexed protein imaging with clickable and cleavable fluorophores

**DOI:** 10.1101/2023.10.20.563323

**Authors:** Thai Pham, Yi Chen, Joshua Labaer, Jia Guo

## Abstract

Single-cell spatial proteomic analysis holds great promise to advance our understanding of the composition, organization, interaction and function of the various cell types in complex biological systems. However, the current multiplexed protein imaging technologies suffer from low detection sensitivity, limited multiplexing capacity or technically demanding. To tackle these issues, here we report the development of a highly sensitive and multiplexed in situ protein profiling method using off-the-shelf antibodies. In this approach, the protein targets are stained with horseradish peroxidase (HRP) conjugated antibodies and cleavable fluorophores via click chemistry. Through reiterative cycles of target staining, fluorescence imaging, and fluoropohore cleavage, many proteins can be profiled in single cells in situ. Applying this approach, we successfully quantified 28 different proteins in a human formalin-fixed paraffin-embedded (FFPE) tonsil tissue, which represents the highest multiplexing capacity among the tyramide signal amplification (TSA) methods. Based on their unique protein expression patterns and their microenvironment, ∼820,000 cells in the tissue are classified into distinct cell clusters. We also explored the cell-cell interactions between these varied cell clusters and observed different subregions of the tissue are composed of cells from specific clusters.

## 1. Introduction

One common feature of complex biological systems, such as brain tissues or solid tumors, is that they are composed of many molecularly and functionally different cell types (1, 2). Single-cell in situ proteomic technologies are powerful tools to study the regulation, function, and interaction of the varied cell types in these heterogeneous biological systems. Mass spectrometry (3) and microarray technologies (4) have wide applications for comprehensive protein analysis. However, as these approaches require the proteins to be isolated from their original cellular context, the spatial information of the proteins is lost during analysis. Immunofluorescence allows proteins to be quantified in their original cellular contexts. However, due to the spectral overlap of the common fluorophores (5), only a handful of proteins can be visualized in one specimen using immunofluorescence.

To enable single-cell spatial proteomic analysis, cyclic immunofluorescence and mass spectrometry imaging have been developed (6). Although these approaches (6-21) allow a large number of proteins to be quantified in their native cellular contexts with the subcellular resolution, some nonideal factors still exist. For instance, in several methods, antibodies are directly labeled with fluorophores or metal isotopes. Without further signal amplification, the detection sensitivity of these technologies can be limited, which hinders their applications to profile low expression proteins or to study specimen with high autofluorescence. Other methods amplify the staining signals using primary antibodies conjugated with oligonucleotides, haptens, or horseradish peroxidase (HRP). Nonetheless, for many protein targets, such chemically modified primary antibodies are not commercially available. And to prepare and validate a panel of those antibodies can be technically demanding, time consuming and expensive.

To enable highly sensitive and multiplexed protein imaging with unconjugated primary antibodies, our laboratory recently developed a spatial proteomic assay using cleavable fluorescent tyramide (CFT) (20). In this approach, protein targets are labeled with off-the-shelf primary antibodies. Subsequently, HRP conjugated secondary antibodies and CFT are applied to stain the targets. After imaging, the fluorophores are chemically cleaved and the antibodies are stripped. Through reiterative cycles, highly sensitive and multiplexed protein imaging can be achieved. HRP catalyzes the conversion of CFT to highly reactive and short-lived radicals, which covalently bind to nearby tyrosine residues. The enzymatic deposition of CFT dramatically increases the signal intensities. The success of this approach requires the molecular weight of the CFT to be relatively small, so that the short-lived radicals can diffuse efficiently and conjugate to many tyrosine residues in close proximity. However, it is highly desirable to have various moieties with large molecular weights to be incorporated into CFT to further improve its performance. For example, the cleavable linkers (22) with different sizes can be applied to enhance signal removal efficiency and reduce the assay time. And quantum dots (23) and polyfluorophores (24, 25) can be used to increase the detection sensitivity and multiplexing capacity.

Here, we report the development of clickable and cleavable fluorophores for ultrasensitive and multiplexed protein imaging. In this approach, instead of depositing cleavable fluorophores directly by HRP, small trans-cyclooctene (TCO) moieties are first conjugated to the tyrosine residues close to the protein targets. Subsequently, click chemistry is applied to couple the cleavable fluorescent tetrazine to CTO by Diels-Alder cycloaddition (26). Using this approach, we successfully labeled the proteins with bulky cleavable fluorophores with multiple cleavage sites. As a result, the signal removal efficiency is significantly improved to 97.5%. Through reiterative cycles of target staining, fluorescence imaging, and fluorophore cleavage, 28 distinct proteins were profiled in single cells of a human formalin-fixed paraffin-embedded (FFPE) tonsil tissue. Based on the unique protein expression patterns and the microenvironment of the individual cells, we partitioned ∼820,000 cells in the same tissue into different clusters. Additionally, we observed that certain cell clusters were predominantly present in specific regions of the tissue, indicating the existence of distinct cellular neighborhoods within the tonsil tissue.

## 2. Material and Methods

### General Information

Chemicals and solvents were obtained from TCI America (Portland, OR, USA) or Sigma-Aldrich (St. Louis, MO, USA) and were directly used without further purification. Bioreagents were purchased from Abcam (Cambridge, United Kingdom), Invitrogen (Waltham, MA, USA), or Novus Biologicals (Littleton, CO, USA), unless otherwise noted.

### Deparaffinization and Antigen Retrieval of FFPE Tonsil Tissue

A total of three xylene deparaffinizations, 10 mintutes each, were performed on tonsil FFPE tissue slides (NBP2-30207, Novus Biologicals, Littleton, Colorado, USA) after heating the slide at 60 °C for 1 hr. The slide was subsequently immersed in 50/50 xylene/ethanol, 100% ethanol, 95% ethanol, and 70% ethanol successively, each for 2 minutes, and then rinsed with deionized water. Afterwards, heat-induced antigen retrieval (HIAR) was performed using a microwave. Antigen retrieval citrate buffer (Abcam ab64236) was applied to the slide and heated for 2 minutes and 45 seconds at high power (700 Watt, level 10) and 14 minutes at low power (140 Watt, level 2) then left to room temperature. To deactivate endogenous horse radish peroxidase (HRP) that may cause false positive signals, the slide was incubated with 3% H_2_O_2_ in PBT (0.1% Triton-X 100 in 1X phosphate buffer saline (PBS)) for 10 minute. Remaining H_2_O_2_ was washed away with PBT twice before proceeding to protein staining.

### Protein staining in FFPE Tissue

To avoid non-specific interaction between the surface of the tissue with staining reagents, the slide was treated with antibody blocking buffer (0.1% (vol/vol) Triton X-100, 1% (wt/vol) bovine serum albumin and 10% (vol/vol) normal goat serum) for 30 minutes at room temperature. The corresponding primary anitbody to the protein of interest (Table 1) was introduced to the slide at the concentration of 5 μg/mL in antibody blocking buffer for 1 hour. The slide was then washed 3 times in PBT. HRP-Conjugated secondary antibody, either goat anti mouse or goat anti rabbit (Table 1) was incubated with the slide for 30 minutes, followed with three times wash with PBT. Afterwards, the slide were stained with TCO-Tyramide at the concentration of 10 nmol/mL in amplification buffer (0.003% H_2_O_2_, 0.1% Tween-20, in 100 mM borate, pH = 8.5) for 10 min at room temperature, and then washed twice with PBT, each for 5 min. Cleavable fluorophore tetrazine (CFTz) was introduced to the slide at the concentration of 5 nmol/mL amplification buffer (100 mM borate, pH = 8.5) for another 10 minutes, and then washed twice with PBT, each for 5 min. To block the uncoupled TCO-Tyramide that might be present in the tissue, 50 nmol/mL of free tetrazine in PBT was added to the tissue and incubated for antoher 10 minutes, and then washed twice with PBT, each for 5 min. The tissues were stained with DAPI and mounted with Prolong Diamond Antifade Reagent before imaging.

**Table 1.** The antibodies used in this study.

| No. | Antibody Target | Catalog No. | Host | Conjugation | Source |
| --- | --- | --- | --- | --- | --- |
| 1 | BRCA1 | 16780 | Mouse | None | Abcam |
| 2 | Ku80 | 232381 | Rabbit | None | Abcam |
| 3 | CD55 | 133684 | Rabbit | None | Abcam |
| 4 | CD79a | 199001 | Mouse | None | Abcam |
| 5 | CD19 | 134114 | Rabbit | None | Abcam |
| 6 | Kat3B | 14984 | Mouse | None | Abcam |
| 7 | CD20 | 279298 | Mouse | None | Abcam |
| 8 | Bcl2 | 182858 | Rabbit | None | Abcam |
| 9 | CD4 | 133616 | Rabbit | None | Abcam |
| 10 | CD45 | 187281 | Mouse | None | Abcam |
| 11 | ILF3 | 206250 | Mouse | HRP | Abcam |
| 12 | NPM1 | 202579 | Mouse | HRP | Abcam |
| 13 | APE1 | 194 | Mouse | None | Abcam |
| 14 | H3S10P | 267372 | Rabbit | None | Abcam |
| 15 | CD11c | 52632 | Rabbit | None | Abcam |
| 16 | H3K9Me2 | 1220 | Mouse | None | Abcam |
| 17 | HLA-DR | 20181 | Mouse | None | Abcam |
| 18 | H3K14Ac | 52946 | Rabbit | None | Abcam |
| 19 | CCR6 | 227036 | Rabbit | None | Abcam |
| 20 | H3K9Ac | 32129 | Rabbit | None | Abcam |
| 21 | H3K18Ac | 40888 | Rabbit | None | Abcam |
| 22 | H3K27Ac | 177178 | Rabbit | None | Abcam |
| 23 | H4K5Ac | 51997 | Rabbit | None | Abcam |
| 24 | H4K8Ac | 45166 | Rabbit | None | Abcam |
| 25 | H4K12Ac | 177793 | Rabbit | None | Abcam |
| 26 | H4K16Ac | 109463 | Rabbit | None | Abcam |
| 27 | Histone H3 | 176842 | Rabbit | None | Abcam |
| 28 | Histone H4 | 177840 | Rabbit | None | Abcam |
| 29 | Rabbit IgG | 6721 | Goat | HRP | Abcam |
| 30 | Mouse IgG | 6789 | Goat | HRP | Abcam |

### Fluorophore Cleavage and HRP Deactivation

100 mM 1,3,5-triaza-7-phosphaadamantane (PTA) and 100 mM tris(2-carboxyethyl)phosphine (TCEP) were successively added to the slide and incubated at 40 °C, each for 30 mintues. After incubation, the slide was taken out and washed threetimes with PBT, each for 5 mintues.

### Antibody Stripping

Antigen retrieval citrate buffer (Abcam ab64236) was added to the slide and heated in the microwave for 2 min and 45 s at high power (level 10, 700 watt) and 14 min at low power (level 2, 140 watt). Then, the slide were submerged in cool water to room temperature for 20 min.

### Syntheis of the Cleavable Fluorescent Tetrazine (CFTz) and tyramide-TCO (Fig. S1-S5)

#### 4-[(1-azido-3-aminopropoxy)methyl]benzoic acid (1)

Methyl 4-[(1-azido-3-trifluoroacetamidopropoxy)methyl]benzoate (2) (0.474 g, 1.3 mmol), synthesized according to the literature(27), was added to a mixture of 4M aqueous sodium hydroxide solution (3.25 mL) and EtOH (3.25 mL). The solution was stirred for 2 hours under room temperature. The solvents were removed by rotary evaporation and the residue was dissolved in 45 mL of D.I. Water. The aqueous layer was extracted with DCM and the organic layer was discarded. The aqueous layer was acidified with 1N HCl to pH 2 and extracted with DCM again. The organic layer was discarded and the aqueous layer was neutralized with 1N NaOH to pH 8. The solution was dried by rotary evaporation and the solids were placed in a funnel with filter paper and washed with 10% MeOH in DCM (45 mL X 2) and 50% MeOH in DCM (45 mL X 2). The solvents were removed by rotary evaporation to obtain the product in the form of mono-sodium salt as a white solid (0.463 g, yield =123.6 %). ^1^H-NMR (500 MHz, MeOD): δ = 1.86-1.90 ppm (m, 2H), 2.69-2.71 ppm (t, 2H), 4.58-4.60 ppm (d, 1H), 4.60-4.63 ppm (t, 1H), 4.77-4.79 ppm(d, 1H), 7.31-7.32 ppm (d, 2H), 7.92-7.93 ppm (d, 2H).

#### N_3_-Cy5 (3)

Cyanine 5 monoacid (1) (10 mg, 0.013 mmol), DMAP (4 mg, 0.034mmol) and DSC (5 mg, 0.034 mmol) were dissolved in anhydrous DMF (0.6 mL) and stirred under room temperature for an hour. TLC (CH_3_OH: CH_2_Cl_2_ = 1:5) was used to check the completion of the reaction. 4-[(1-azido-3-aminopropoxy) methyl] benzoic acid (1) (9.6 mg, 0.034 mmol) and DIPEA (7 μL, 0.041 mmol) was added and the solution was stirred for another hour. The solvent was removed by rotary evaporation and the crude was further purified through preparative silica gel TLC plate (25 X 25 cm; silica gel 60; CH_3_OH: CH_2_Cl_2_ = 1:5). Dark blue solid was obtained as the product.

#### Tetrazine-N_3_-Cy5 (5)

N_3_-Cy5 (3) (11.8 mg, 0.014 mmol), TSTU (6.1 mg, 0.02 mmol) and DIPEA (3.5 μL, 0.02 mmol) was dissolved in anhydrous DMF (0.6mL) and stirred for an hour at room temperature. TLC (CH_3_OH: CH_2_Cl_2_ = 1:5) was used to check the completion of the reaction. Methyl tetrazine amine (4 mg, 0.02 mmol) and DIPEA (3.5 μL, 0.02 mmol) was added and the solution was stirred for another hour. The solvent was removed by rotary evaporation and the crude was further purified through preparative preparative silica gel TLC plate (25 X 25 cm; silica gel 60; CH_3_OH: CH_2_Cl_2_ = 1:5) to obtain a dark blue solid. The residue was further purified by semi-preparative reverse phase HPLC HPLC gradient: A, 100% 0.1 M TEAA; B 100% MeCN; 0-2 min, 5% B (flow 2-5 ml/min); 2-10 min, 5-22% B (flow 5 ml/min); 10-15 min, 22-30% B (flow 5 ml/min); 15-20 min, 30-40% B (flow 5 ml/min); 20-25 min, 40-50% B (flow 5 ml/min); 25-30 min, 50-60% B (flow 5 ml/min); 30-32 min, 60-70% B (flow 5 ml/min); 32-35 min, 70-95% B (flow 5 ml/min); 35-37 min, 95% B (flow 5ml/min); 37-39 min, 95-5% B, (flow 5 ml/min); 39-42 min, 5% B (flow 5-2 ml/min)]. The fraction with retention time of 26.40 min was collected and evaporated under reduced pressure to afford Tetrazine-N_3_-Cy5 (5) (4.22 mg) as a dark blue solid. ^1^H-NMR (500MHz, MeOD): δ = 1.40-1.43 ppm (t, 6H), 1.63-1.66 ppm (t, 2H), 1.77ppm (s,12H), 2.11-2.17 ppm (t, 2H), 3.05 ppm (s, 3H), 4.06-4.09 ppm (t, 2H), 4.17-4.21 ppm (q, 2H), 4.68-4.70 ppm (m, 2H), 4.73 ppm (s, 2H), 4.92 ppm (s, 1H), 6.32-6.38 ppm (t, 2H), 6.68-6.73 ppm (t, 1H), 7.31-7.38 ppm (dd, 2H), 7.53-7.55 (d, 2H), 7.65-7.66 (d, 2H), 7.89-7.95 ppm (m, 6H), 8.30-8.35 ppm (t, 2H), 8.53-8.55 ppm (d, 2H). HRMS [+ Scan]; calculated m/z for C_54_H_62_N_11_O_9_S_2_^+^: 1070.4022; observed m/z: 1070.3948.

#### N_3_-N_3_-Cy5 (6)

N_3_-Cy5 (3) (10.5 mg, 0.012 mmol) (Scheme 5.2), TSTU (6.1 mg, 0.02 mmol) and DIPEA (3.5 μL, 0.02 mmol) was dissolved in anhydrous DMF 0.6mL and stirred under room temperature for an hour. TLC (CH_3_OH: CH_2_Cl_2_ = 1:5) was used to check the completion of the reaction. 4-[(1-azido-3-aminopropoxy) methyl] benzoic acid (1) (4.8 mg, 0.017 mmol) and DIPEA (3.5 μL, 0.02 mmol) was added and the solution was stirred for another hour. The solvent was removed by rotary evaporation and the crude was further purified through preparative silica gel TLC plate (25 X 25 cm; silica gel 60; CH3OH: CH2Cl2 = 1:5). The product was obtained as a dark blue solid.

#### Tetrazine-N_3_-N_3_-Cy5 (7)

N_3_-N_3_-Cy5 (6) (11.1 mg, 0.01 mmol), TSTU (6.1 mg, 0.02 mmol) and DIPEA (3.5 μL, 0.02 mmol) was dissolved in anhydrous DMF 1mL and stirred under room temperature for an hour. TLC (CH_3_OH: CH_2_Cl_2_ = 1:5) was used to check the completion of the reaction. Methyl tetrazine amine (2.5 mg, 0.012 mmol) and DIPEA (3.5 μL, 0.02 mmol) was added and the solution was stirred for anotherhour. The solvent was removed by rotary evaporation and the crude was further purified through preparative TLC (CH_3_OH: CH_2_Cl_2_= 1: 5). The product was obtained as a dark blue solid. The residue was further purified by semi-preparative reverse phase HPLC HPLC gradient: A, 100% 0.1 M TEAA; B 100% MeCN; 0-2 min, 5% B (flow 2-5 ml/min); 2-10 min, 5-22% B (flow 5 ml/min); 10-15 min, 22-30% B (flow 5 ml/min); 15-20 min, 30-40% B (flow 5 ml/min); 20-25 min, 40-50% B (flow 5 ml/min); 25-30 min, 50-60% B (flow 5 ml/min); 30-32 min, 60-70% B (flow 5 ml/min); 32-35 min, 70-95% B (flow 5 ml/min); 35-37 min, 95% B (flow 5ml/min); 37-39 min, 95-5% B, (flow 5 ml/min); 39-42 min, 5% B (flow 5-2 ml/min)]. The fraction with retention time of 28.35 min was collected and evaporated under reduced pressure to afford Tetrazine-N_3_-N_3_-Cy5 (7) (3.21 mg) as a dark blue solid. ^1^H-NMR (500MHz, MeOD): δ = 0.86-0.89 ppm (t, 6H), 1.48-1.57 ppm (t, 4H), 1.76 ppm (s,12H), 2.15 -2.196 ppm (t, 4H), 3.06 ppm (s, 3H), 3.36-3.43 ppm (t, 2H), 4.25-4.27 ppm (q, 2H), 4.78-4.80 ppm (t, 2H), 4.84 ppm (s, 1H), 4.94ppm (s, 1H), 6.33-6.38 ppm (t, 2H), 6.68-6.72 ppm (t, 1H), 7.33-7.38 ppm (dd, 2H), 7.44-7.45 (d, 2H), 7.31-7.53 (d, 2H), 7.66-7.67 ppm (q, 4H), 47.75-7.77 ppm (q, 4H), 8.30-8.37 ppm (t, 2H), 8.55-8.57 ppm (d, 2H). HRMS [+ Scan]; calculated m/z for C_65_H_74_N_15_O_11_S_2_^+^: 1302.4983; observed m/z: 1302.4899.

#### N_3_-N_3_-N_3_-Cy5 (9)

N_3_-N_3_-Cy5 (6) (16.0 mg, 0.14 mmol), TSTU (9.2 mg, 0.03 mmol) and DIPEA (5.2 μL, 0.03 mmol) was dissolved in anhydrous DMF 0.6 mL and stirred under room temperature for an hour. TLC (CH_3_OH: CH_2_Cl_2_ = 1:5) was used to check the completion of the reaction. 4-[(1-azido-3-aminopropoxy) methyl] benzoic acid (1) (5.6 mg, 0.02 mmol) and DIPEA (3.5 μL, 0.02 mmol) was added and the solution was stirred for another hour. The solvent was removed by rotary evaporation and the crude was further purified through preparative silica gel TLC plate (25 X 25 cm; silica gel 60; CH_3_OH: CH_2_Cl_2_ = 1:5). The product was obtained as a dark blue solid.

#### Tetrazine-N_3_-N_3_-N_3_-Cy5 (11)

N_3_-N_3_-N_3_-Cy5 (9) (13.5 mg, 0.01 mmol), TSTU (6.1 mg, 0.02 mmol) and DIPEA (3.5 μL, 0.02 mmol) was dissolved in anhydrous DMF 1mL and stirred under room temperature for an hour. TLC (CH_3_OH: CH_2_Cl_2_ = 1:5) was used to check the completion of the reaction. DIPEA (3.5 μL, 0.02 mmol) and methyl tetrazine amine (2.5 mg, 0.012 mmol) were added and the mixed solution was stirred for another hour. The solvent was removed by rotary evaporation and the crude was further purified through preparative TLC (CH_3_OH: CH_2_Cl_2_ = 1:5). The product was obtained as a dark blue solid. The residue was further purified by semi-preparative reverse phase HPLC HPLC gradient: A, 100% 0.1 M TEAA; B 100% MeCN; 0-2 min, 5% B (flow 2-5 ml/min); 2-10 min, 5-22% B (flow 5 ml/min); 10-15 min, 22-30% B (flow 5 ml/min); 15-20 min, 30-40% B (flow 5 ml/min); 20-25 min, 40-50% B (flow 5 ml/min); 25-30 min, 50-60% B (flow 5 ml/min); 30-32 min, 60-70% B (flow 5 ml/min); 32-35 min, 70-95% B (flow 5 ml/min); 35-37 min, 95% B (flow 5ml/min); 37-39 min, 95-5% B, (flow 5 ml/min); 39-42 min, 5% B (flow 5-2 ml/min)]. The fraction with retention time of 33.8 min was collected and evaporated under reduced pressure to afford Tetrazine-N_3_-N_3_-N_3_-Cy5 (11) (1.55 mg) as a dark blue solid. ^1^H-NMR (500MHz, MeOD): δ = 0.86-0.89 ppm (t, 6H), 1.61-1.68 ppm (t, 6H), 1.76 ppm (s,12H), 2.08-2.19 ppm (m, 6H), 3.06 ppm (s, 3H), 3.46-3.60 ppm (t, 2H), 4.14-4.21 ppm (q, 2H), 4.62-4.68 ppm (t, 1H), 4.70 ppm (s, 2H), 4.72-4.75 ppm (t, 1H), 4.76-4.79 ppm (t,1H), 4.83 ppm (s, 1H), 4.91-4.92 ppm (s,1H), 6.35-6.38 ppm (t, 2H), 6.67-6.73 ppm (t, 1H), 7.32-7.36 ppm (dd, 2H), 7.42-7.44 (d, 2H), 7.51-7.53 ppm (d,2H) 7.63-7.64 ppm (d, 2H), 7.70-7.72 (d, 2H), 7.74-7.75 ppm (d, 2H), 7.89-7.93 ppm (m, 6H), 8.33-8.35 ppm (t, 2H), 8.54-8.55 ppm (d, 2H). HRMS [+ Scan]; calculated m/z for C_76_H_86_N_19_O_13_S_2_^+^: 1534.5943; observed m/z: 1534.6039.

#### Tyramide-TCO (12)

TCO-(PEG)_4_-NHS ester (13) (6.0 mg, 0.012 mmol) and Tyramine (14) (2.0 mg, 0.011 mmol) were diluted in 10 μL anhydrous DMF. The mixture was kept in dark room for 2 hours to form the product. The solution was then further diluted in the total of 0.6 mL of DMF as the stock solution for IHC staining.

#### Imaging and Data Analysis

A 20x obejective equiped Nikon epifluoresecent microscope was wont to image the FFPE tissue. Tissue image was captured by a CooSNAP HQ2 camera and C-FL DAPI HC HISN via Chroma 49009 filter. NIS-Element Imaging Software was used to process the obtained image data. To allign all the staining images, DAPI image from each cycle was used as the coordination reference. To generate single cell protein expression profile, cells are defined based on nuclear DAPI staining using NIS Elements Imaging software. Regions of interest (ROIs) were determined by expanding the DAPI signal on every single cell by 10 pixels. Signal intensity values within these ROIs were then calculated via Cell Profiler resulting in a comma separated value (CSV) files. These files were then unsupervisedly clustered to generate Optsne plot (https://doi.org/10.1038/s41467-019-13055-y) pseudo-color images were generated with ImageJ. Cell neighborhoods were calculated by detecting and classifying the encompassing cells within 20 μm or less of every individual cell in the sample. The quantity of cells from the various clusters in each cell neighborhood were used for clustering to come up with subcluster Optsne plots.

## 3. Results

### Platform design

In this ultrasensitive and multiplexed protein imaging method, each analysis cycle consists of six major steps, as shown **Figure 1**. First, off-the-shelf primary antibodies from different species or of varied immunoglobulin classes are applied to bind to the protein targets. Second, one protein target is then stained with HRP conjugated primary or secondary antibodies and fluorescently labeled. Third, images are captured under a fluorescence microscope, generating single-cell protein expression profiles. To facilitate image alignment, the nucleus is also stained with DAPI and imaged along with the protein target. Fourth, the fluorophores are cleaved chemically, and HRP is simultaneously deactivated. Fifth, steps two to four are repeated until all protein targets in this analysis cycle are detected. Finally, all antibodies are stripped to initiate the next analysis cycle. By repeating cycles of protein staining, fluorescence imaging, fluorophore removal, HRP deactivation, and antibody stripping, this technology enables ultrasensitive and multiplexed in situ protein profiling in single cells of intact tissues.

**Figure 1:**
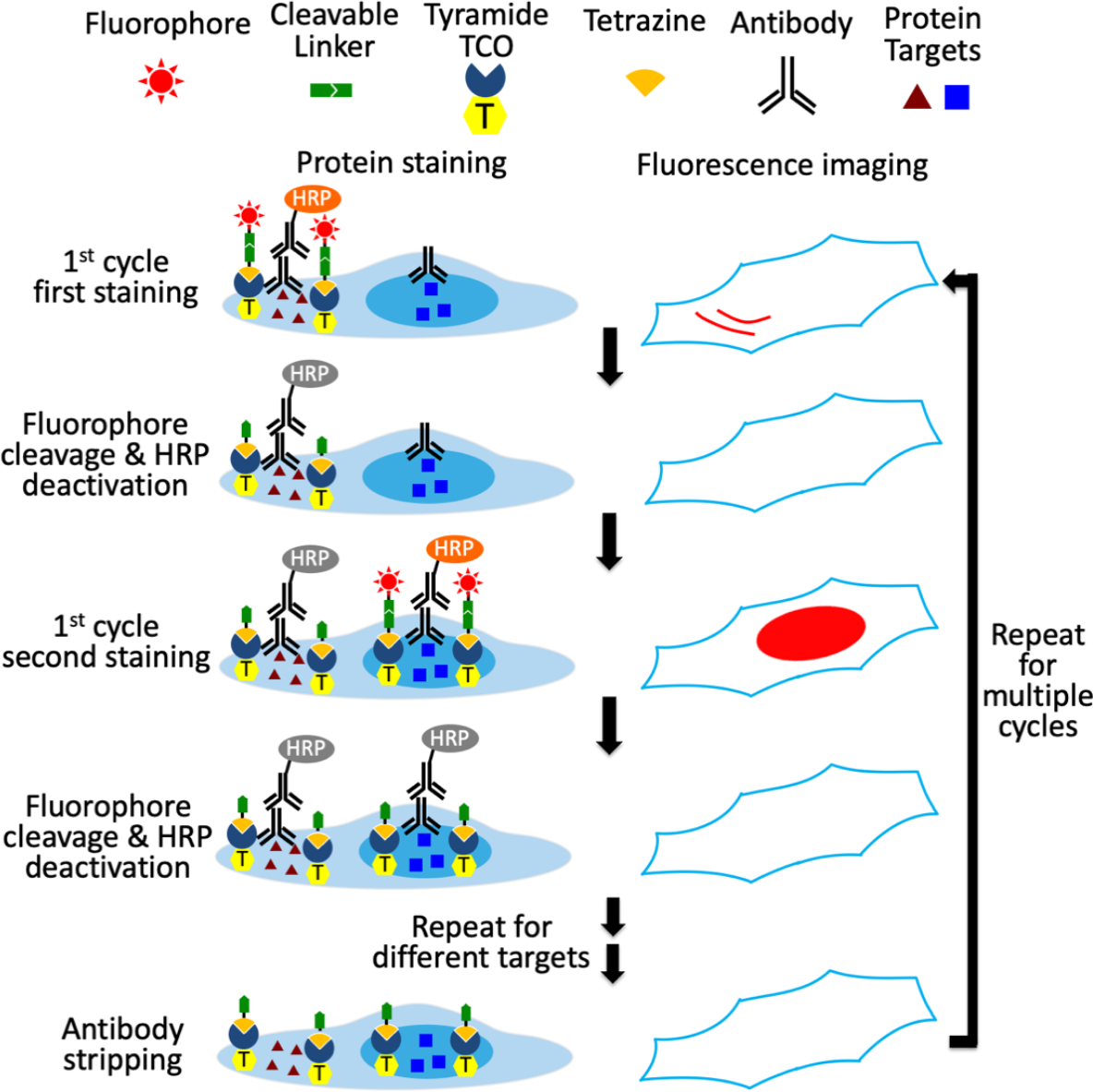
Ultrasensitive and multiplexed protein imaging with cleavable fluorescent tetrazine (CFTz). In each cycle, multiple protein targets are first recognized by different primary antibodies. Subsequently, the first target is stained by HRP-conjugated antibodies and TCO-tyramide. The cleavable fluorecent tetrazine (CFTz) are then covalently coupled to the TCO-tyramide via click chemistry reaction to stain the target protein. After imaging, the fluorophores are chemically cleaved and HRP is simultaneously deactivated. The processes of protein staining, fluorescence imaging, fluorophore cleavage and HRP deactivation are repeated until every target in the first cycle is stained. Finally, all the antibodies are stripped to initiate the next cycle. Through reiterative analysis cycles, a large number of distinct proteins can be quantitatively profiled in single cells in situ.

To minimize the size of HRP substrates while ensuring the large moieties to be successfully applied for target staining, we split the staining probes into two compartments (**Figure 2**). The first compartment is consisted of the tyramide moiety and trans-cyclooctene (TCO); while the second compartment contains the fluorophores tethered to tetrazine through cleavable linkers. The small size of tyramide-TCO allows it to be efficiently deposited by HRP to conjugate with many tyrosine residues proximal to the protein of interest. Subsequently, cleavable fluorescent tetrazine (CFTz) are coupled to the deposited TCO moieties through an irreversible and bioorthogonal Diels Alders cycloaddition. Unlike the direct labeling with short-lived (<1 ms) radicals, this click chemistry reaction enables much longer reaction time (mins to hrs). As a result, the relatively large-sized CFTz can be successfully applied for target staining. To assess the effectiveness of this approach, we designed CFTz with one (Tetrazine-N_3_-Cy5), two (Tetrazine-N_3_-N_3_-Cy5) or three (Tetrazine-N_3_-N_3_-N_3_-Cy5) cleavage sites. As long as one site is cleaved, the fluorescent signals could be successfully erased. Thus, the CFTz with multiple cleavage sites should have higher signal removal efficiency, which could lead to improved multiplexing capacity and analysis accuracy.

**Figure 2.**
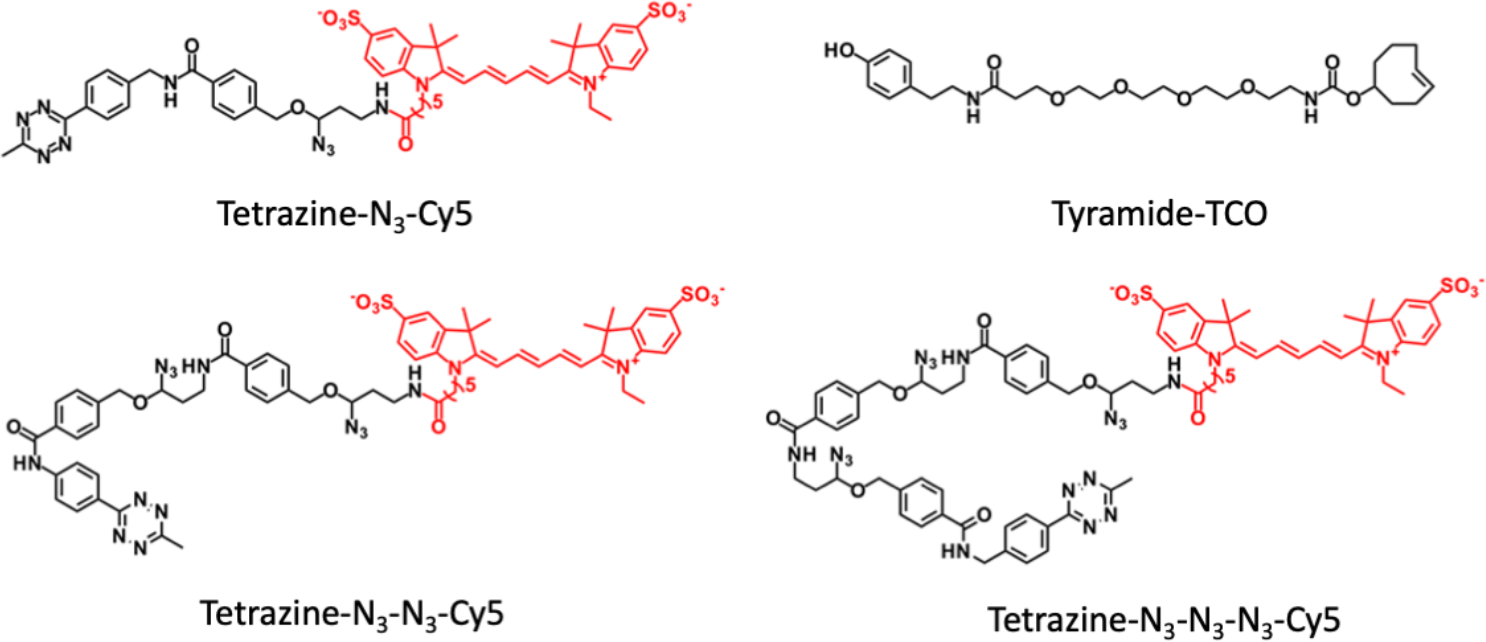
Chemical structures of CFTz (Tetrazine-N_3_-Cy5, Tetrazine-N_3_-N_3_-Cy5, and Tetrazine-N_3_-N_3_-N_3_-Cy5) and Tyramide-CTO.

### Efficient staining by CFTz with multiple cleavage sites

To evaluate whether proteins can be effectively stained by clickable and cleavable CFTz, we labeled protein CD45 in human tonsil FFPE tissues. Due to the fast reaction kinetics of the click chemistry, 1-linker and 2-linker CFTz reached ∼90% and ∼80% of the maximum fluorescence intensity within 10 min, respectively. For 3-linker CFTz, it requires 30 min to obtain ∼80% of the maximum fluorescence intensity (Figure 3A). The staining patterns obtained with CFTz are consistent with the conventional tyramide signal amplification (TSA) assay (Figure 3B). And the staining intensities generated by TSA, 1-linker and 2-linker CFTz closely resemble each other; while the intensities obtained by 3-linker CFTz is ∼25% lower (Figure 3C). These results indicate proteins in FFPE tissues can be efficiently stained by CFTz with a short labeling time.

**Figure 3.**
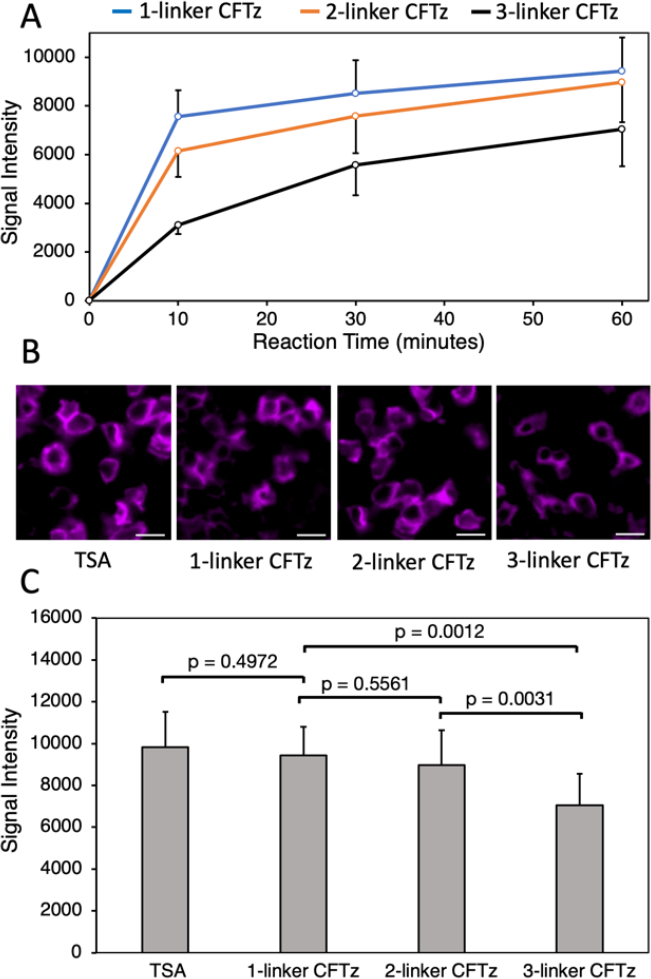
(A) The signal intensities obtained by staining protein CD45 with 1-linker, 2-linker, and 3-linker CFTz in human tonsil FFPE tissues for 10, 30 and 60 min, respectively. (B) Protein CD45 in human tonsil FFPE tissues is stained by conventional tyramide signal amplification (TSA), 1-linker, 2-linker, or 3-linker CFTz. Scale bars, 10 μm. (C) Comparison of the staining intensities generated by TSA, 1-linker, 2-linker, and 3-linker CFTz.

### Enhanced cleavage efficiency by CFTz with multiple cleavage sites

To assess the fluorophore cleavage efficiency of CFTz, we stained the protein CD45 and subsequently treated the slides with the mild reducing reagents tris (2-carboxyethyl) phosphine (TCEP) and1,3,5-Triaza-7-phosphaadamantane (PTA) (Figure 4A). After cleavage, almost all the fluorescence signals are removed (Figure 4B). We then quantified and compared the cleavage efficiency of 1-linker, 2-linker, and 3-linker CFTz (Figure 4C). 1-linker CFTz has the cleavage efficiency of ∼95.5%, which is consistent with our previously developed cleavable fluorescent probes (12, 17-20). With multiple cleavage sites, 2-linker, and 3-linker CFTz increased the cleavage efficiency to ∼97.5%. By reducing the signal leftover by ∼50% in each analysis cycle, CFTz with multiple cleavage sites will significantly enhance the protein quantification accuracy and the multiplexing capacity of reiterative protein staining approaches.

**Figure 4:**
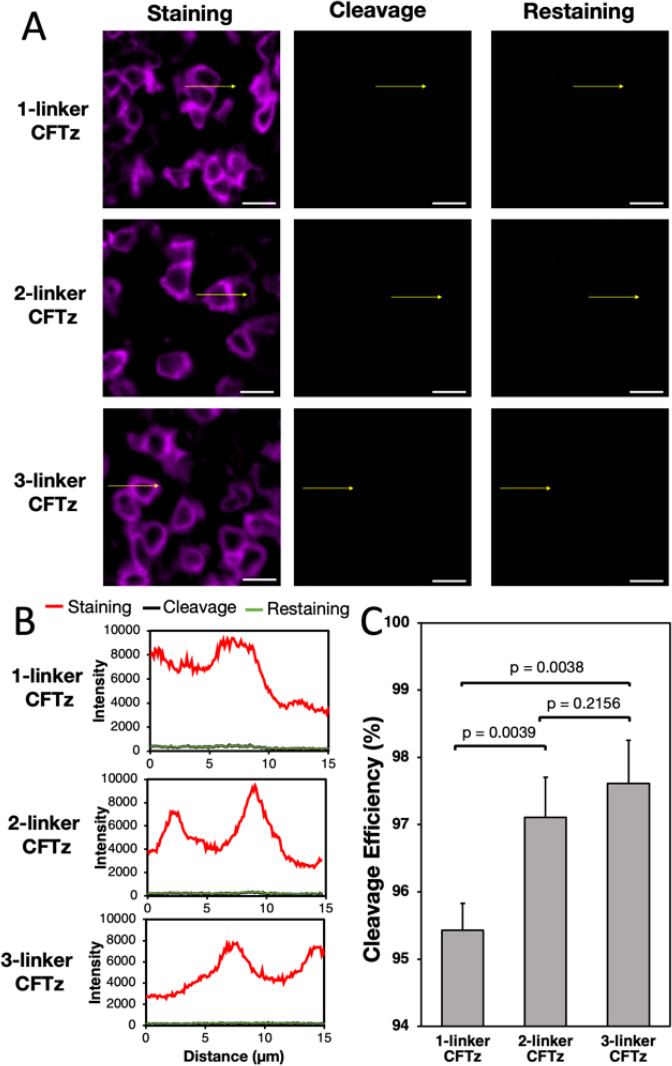
(A) Protein CD45 is stained with 1-linker, 2-linker, and 3-linker CFTz in human tonsil FFPE tissues (left). Subsequently, the fluorescence signals are removed by mild reducing reagents (middle). Afterwards, the same tissues are blocked with free tetrazine and restained with CFTz (right). Scale bars, 10 μm. (B) Fluorescence intensity profiles corresponding to the indicated arrow positions in (A). (C) Comparison of the cleavage efficiencies for 1 linker, 2 linker, and 3 linker CFTz.

To ensure that all the unreacted TCO are quenched, we incubated the tissues with free tetrazine following cleavage. Then, we restained the tissues with CFTz, and observed no fluorescence signal increase (Figure 4A and 4B). These results suggest that the unreacted TCO are efficiently quenched at the end of each staining cycle. Thus, it will not lead to false positive signals in subsequent protein quantification. Additionally, as documented in our previous studies, the TCEP, PTA and antibody stripping treatments will not damage the integrity of epitopes (19, 20). Therefore, the CFTz with the improved cleavage efficiency should enable a large number of varied proteins to be accurately quantified in single cells in situ.

### Multiplexed In situ protein profiling in the Human Tonsil Tissue

To demonstrate the feasibility of this approach for multiplexed protein imaging, we stained 28 different proteins in 28 sequential analysis cycles on a human FFPE tonsil tissue. Due to its fast reaction kinetics and high cleavage efficiency, 2-linker CFTz (Tetrazine-N_3_-N_3_-Cy5) was applied to carry out the study. Using off-the-shelf antibodies, all the 28 proteins were successfully detected with the subcellular resolution (Figure 5). The obtained staining patterns are consistent with the ones generated by conventional immunohistochemistry (28). With the enzymatic signal amplification by HRP, our approach preserves the super-high sensitivity of the TSA assay. Thus, our approach enables proteins to be unambiguously detected in highly autofluorescent FFPE tissues with millisecond exposure time and a 4x objective. This setup allows 1-2 cm^2^ tissue to be imaged within 5 min, which dramatically reduces the assay time and enhances the sample throughput. Additionally, by improving the cleavage efficiency by CFTz with multiple cleavage sites, our method allows more proteins to be precisely quantified in the same specimen. By quantifying 28 proteins in the same tissue, our approach has the highest multiplexing capacity among all the TSA based assays.

**Figure 5.**
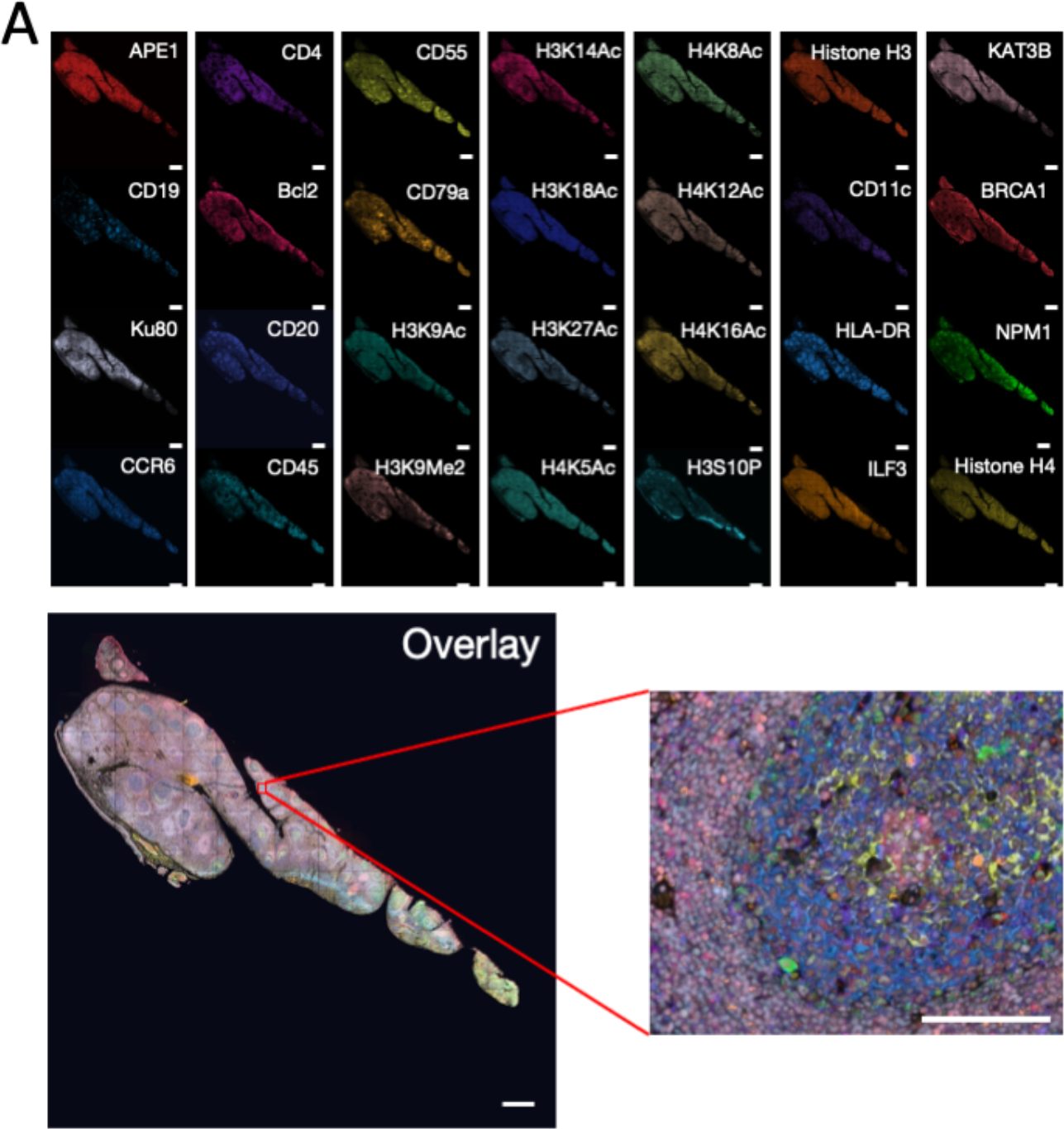
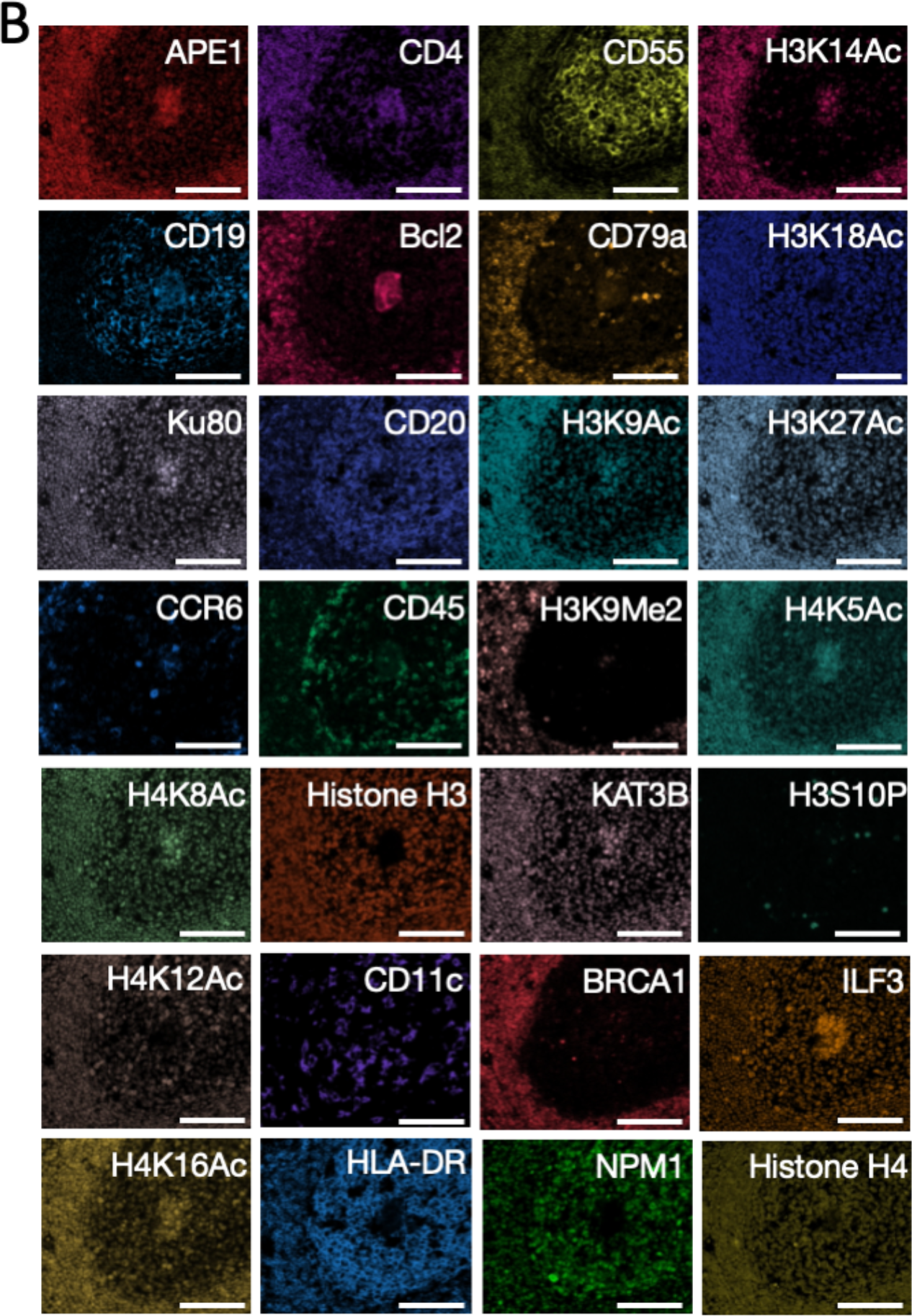
(A) 28 diferent proteins are stained with CFTz on the same FFPE Tissue. Scale bar, 500μm. (B):Zooned view in the boxed area in (A). Scale bar, 100μm

### Different Cell Types and Their Spatial Distribution in the Human Tonsil Tissue

We utilized the single-cell in situ protein expression profiles to investigate the heterogeneity and spatial distribution of various cell types in the human tonsil tissue. To accomplish this, we calculated the expression levels of the 28 proteins in each of the ∼820,000 cells identified in the tissue. Based on their unique protein expression patterns (Figure 6A and Supplementary Figure S9), we partitioned the cells into 14 distinct cell clusters (Figure 6B) using the Optsne software (29). These cell clusters were then mapped back to their natural tissue locations (Figure 6C and Supplementary Figure S10), revealing that different subregions of the tonsil tissue were predominantly composed of cells from specific clusters. For example, cluster 4 was mainly present in the epithelium; clusters 2 and 3 dominated the submucosa and connective capsule; clusters 1 and 5 were the major cell type in the lymphoid follicles; clusters 8, 9, 11, 13 and 14 dominated the germinal centers; and clusters 6, 7 and 12 were exclusive to the connective tissues. These results demonstrate that our approach enables the classification of cell types and study of their spatial distribution in formalin-fixed paraffin-embedded (FFPE) tissues.

**Figure 6.**
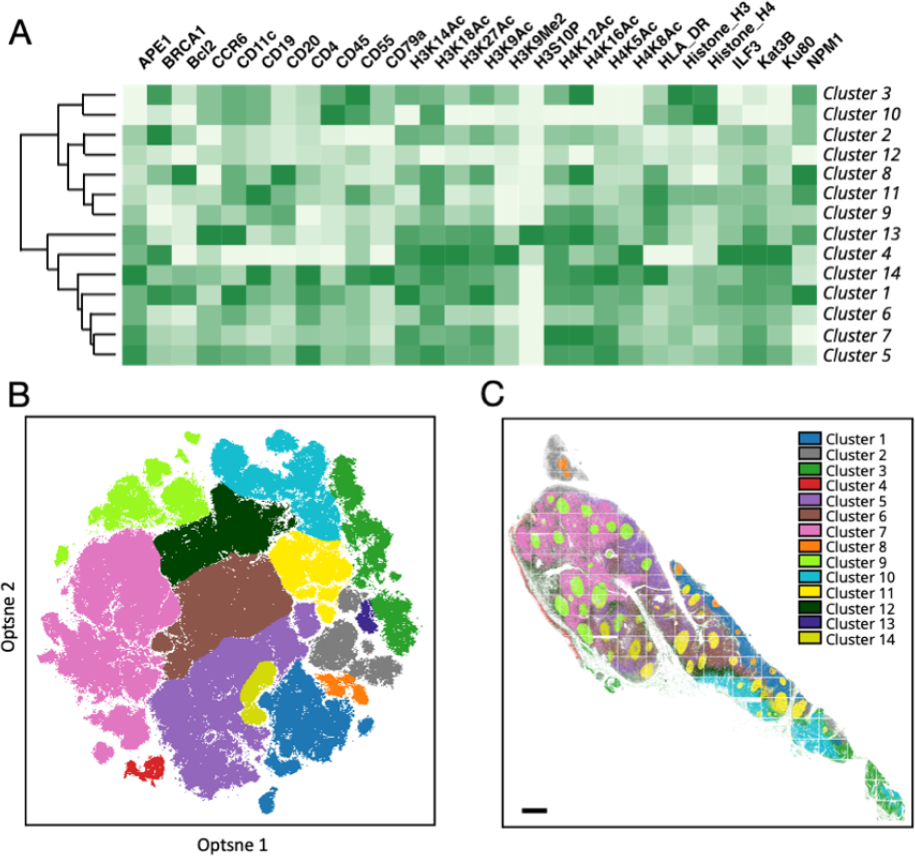
(A) Based on their different single cell protein expression profiles, (B) ∼820,000 individual cells in the human tonsil tissue are partitioned into 14 cell clusters. (C) Anatomical locations of each cell from the 14 clusters. Scale bar, 500 μm.

### Cell-Cell Interaction in the Human Tonsil Tissue

By classifying the cell types in their native spatial contexts, our approach also enables the exploration of the cell-cell interactions among the various cell clusters (Figure 7). The term “cell neighborhood” was defined as all the cells within the distance of 20 μm from the center cell. For each single cell in the tissue, we counted the number of cells from different clusters in its neighborhood. Then, we calculated the averaged cell numbers of each cluster in varied cell neighborhoods. By projecting the metrics on a heatmap, we observed a significant association between cell clusters 5 and 6, 9 and 13, and 5 and 14. Interestingly, we found a consistently strong association between cells from the same cell cluster (Figure 7, diagonal), while most of the cells from distinct clusters avoided contact. These results suggest that homotypic cell adhesion may play a crucial role in shaping the architecture of human tonsil tissue.

**Figure 7.**
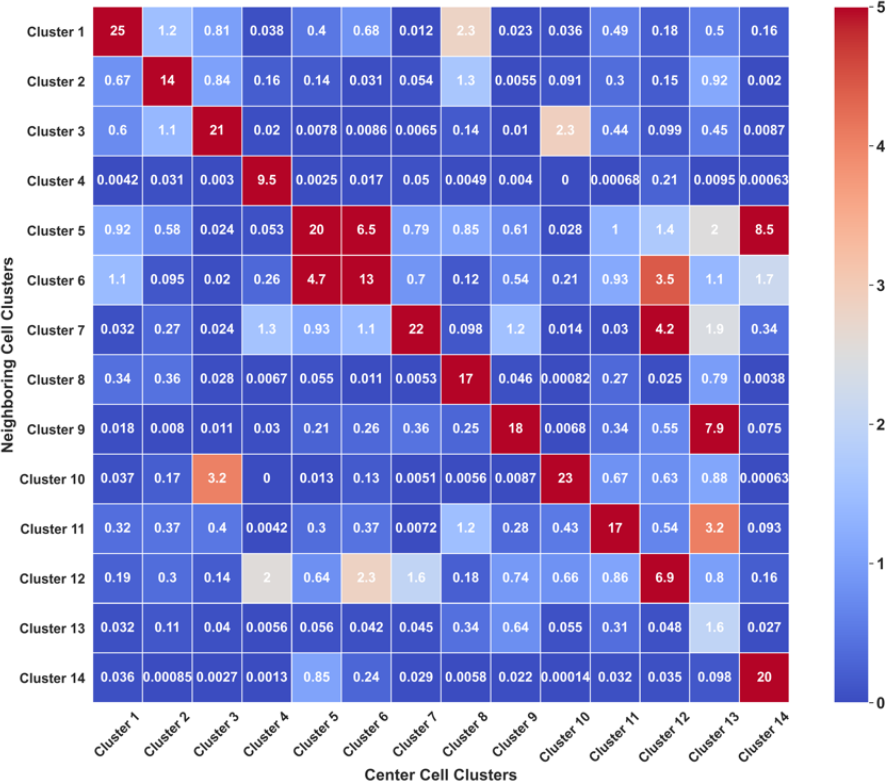
Averaged cell numbers of each cell cluster in different cell neighborhoods.

The individual cells from the same cluster can also be further classified into subclusters, based on the cells in their neighborhood. By mapping these subclusters back to their original tissue locations, we observed that varied subclusters from the same cell cluster are located in specific subregions of the tonsil tissue. For instance, cluster 13 was further partitioned into 2 subclusters: 13a and 13b (Figure 8A and 8B).

**Figure 8.**
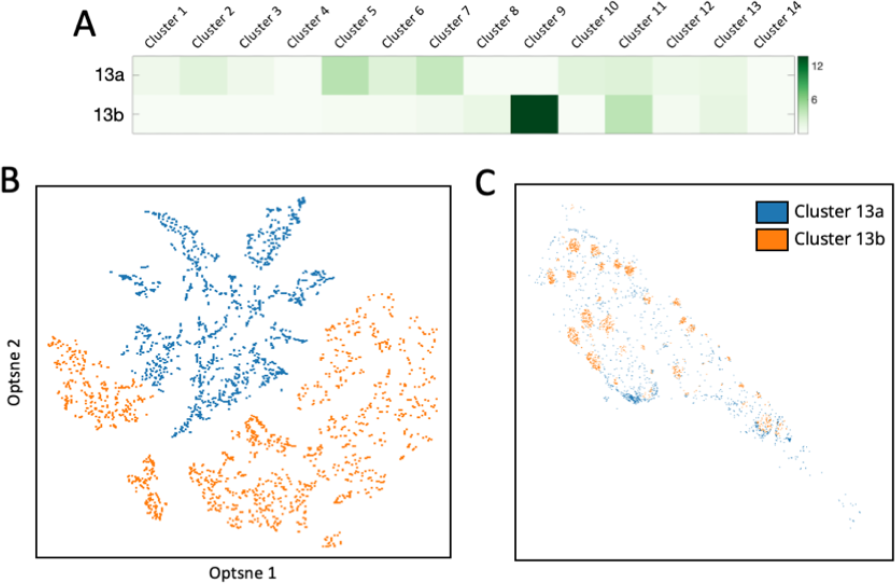
(A) Based on their neighbor cells from different clusters, (B) the individual cells in cluster 13 are further partitioned into 2 subclusters. (C) Anatomical locations of each cell from the three subclusters

Subcluster 13a was predominantly present in the germinal centers; while subcluster 13b was mainly in connective tissues (figure 8C). These findings suggest our approach enables the studies of cell-cell interaction and cell type classification based on their microenvironment.

## 4. Discussion

In this study, we have successfully developed clickable and cleavable fluorescent probes, and applied them for highly sensitive and multiplexed single-cell in-situ protein profiling. Our approach has overcome the limitation that only the small probes can be applied in the TSA assay. Thus, the bulky fluorophores, such as quantum dots (23) and polyfluorophores (24, 25), can also be applied for HRP-catalyzed enzymatic target staining. Using this approach, we have designed and synthesized the fluorescent tyramide with multiple cleavage sites. With these probes, the signal removal efficiency is significantly improved, compared to probes with only one cleavage site. As a result, our approach enhances the analysis accuracy and the multiplexing capacity of the existing multiplexed protein imaging technologies.

With our method, we have demonstrated the ability to classify individual cells in human tonsil tissue into distinct cell clusters using their unique multiplexed protein expression profiles. Furthermore, our analysis revealed that different subregions of the tonsil tissue are composed of cells originating from diverse clusters. We also investigated the associations and avoidance patterns between various cell clusters. Additionally, we observed that the single cells within each cluster can be further partitioned into subclusters based on the distinctive cell clusters of their neighboring cells. These findings highlight the potential of our approach to classify cell types and subtypes by leveraging protein expression profiles and the characteristics of neighboring cells. The identification of these cell types and subtypes holds promise for advancing our understanding of cell heterogeneity, facilitating disease diagnosis, and enabling patient stratification.

The number of imaging cycles and the number of proteins quantified in each cycle are the key factors determining the multiplexing capacity of our approach. We have previously reported that our signal erasing treatment and antibody stripping process did not cause any damages to the integrity of the epitopes (19, 20). Here, we have demonstrated that at least 28 reiterative cycles can be carried out on the same tissue. In each analysis cycle, varied protein targets can be labeled simultaneously with antibodies from varied species or of different immunoglobin classes, hapten or HRP-conjugated primary antibodies. By employing CFTz with four or five distinct fluorophores, along with iterative protein staining, HRP deactivation and fluorophore cleavage, potentially over 10 proteins can be imaged before antibody stripping in every analysis cycle. Consequently, we anticipate that this multiplexed protein imaging technique holds the potential to profile numerous protein targets, potentially in the hundreds, within the same specimen.

The clickable and cleavable fluorescent probes developed here can also be applied for highly sensitive and multiplexed nucleic acid (30-38) and metabolic (39) imaging. The integration of these technologies and our approach will enable the comprehensive in situ profiling of DNA, RNA, proteins, and metabolites at the single-cell level within intact tissues. Additionally, the incorporation of a program-controlled microfluidic system (40) with a standard fluorescence microscope will make an automated tissue imaging platform. The combination of these advancements forms a highly multiplexed molecular imaging system with broad applications in systems biology and biomedical research.

## Data Availability

All the data is included in the manuscript and supporting information.

