## Supplementary material for "Ultrasensitive and multiplexed protein imaging with clickable and cleavable fluorophores": SI figures

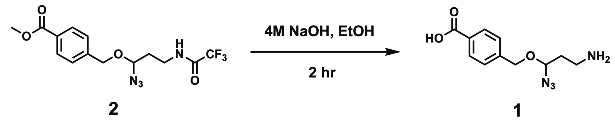


Figure S1. Synthetic Scheme of azido cleavable linker

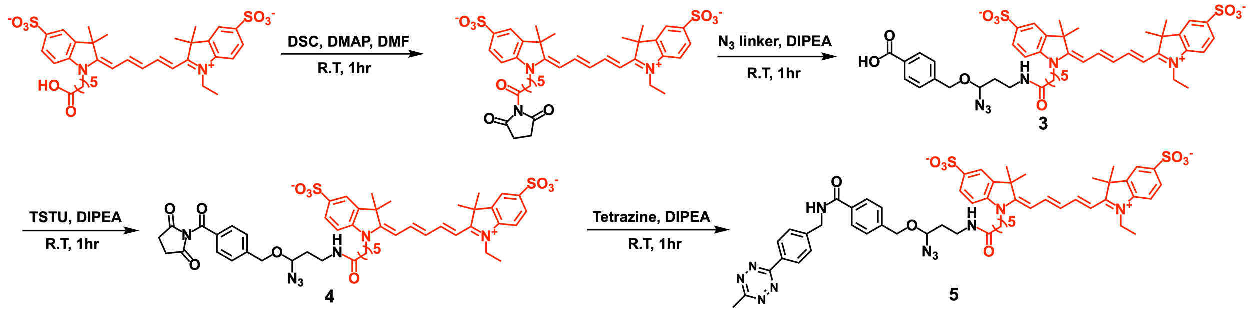


Figure S2. Synthetic Scheme of Tetrazine-N3-Cy5

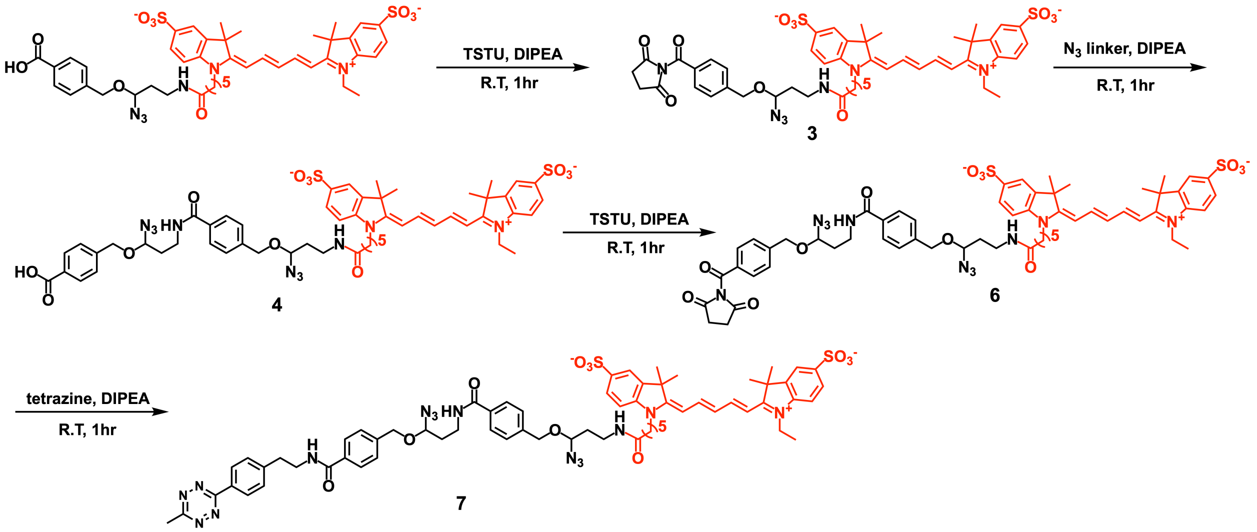


Figure S3. Synthetic Scheme of Tetrazine-$N_3$-$N_3$-Cy5

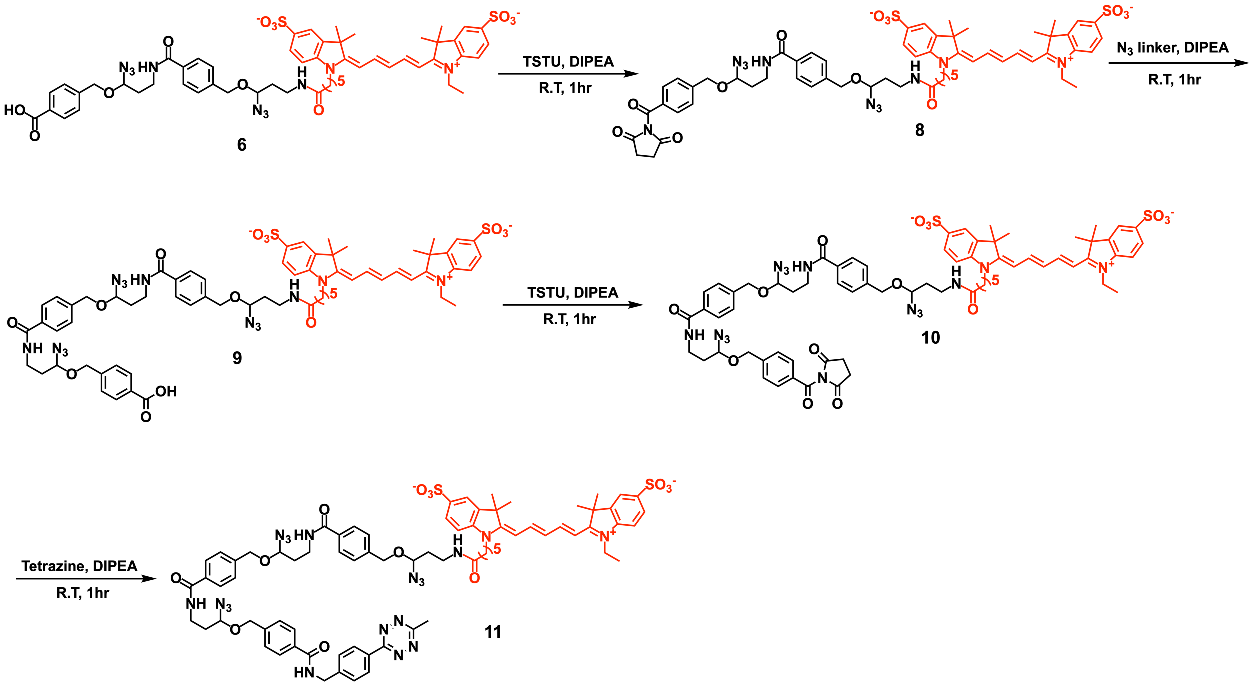


Figure S4. Synthetic Scheme of Tetrazine-$N_3$-$N_3$-$N_3$-Cy5

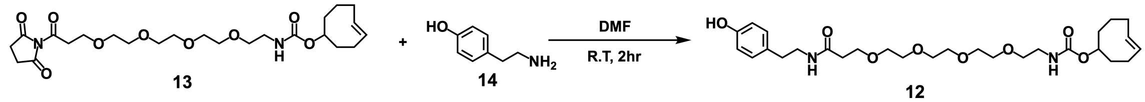


Figure S5. Synthetic Scheme of Tyramide-TCO

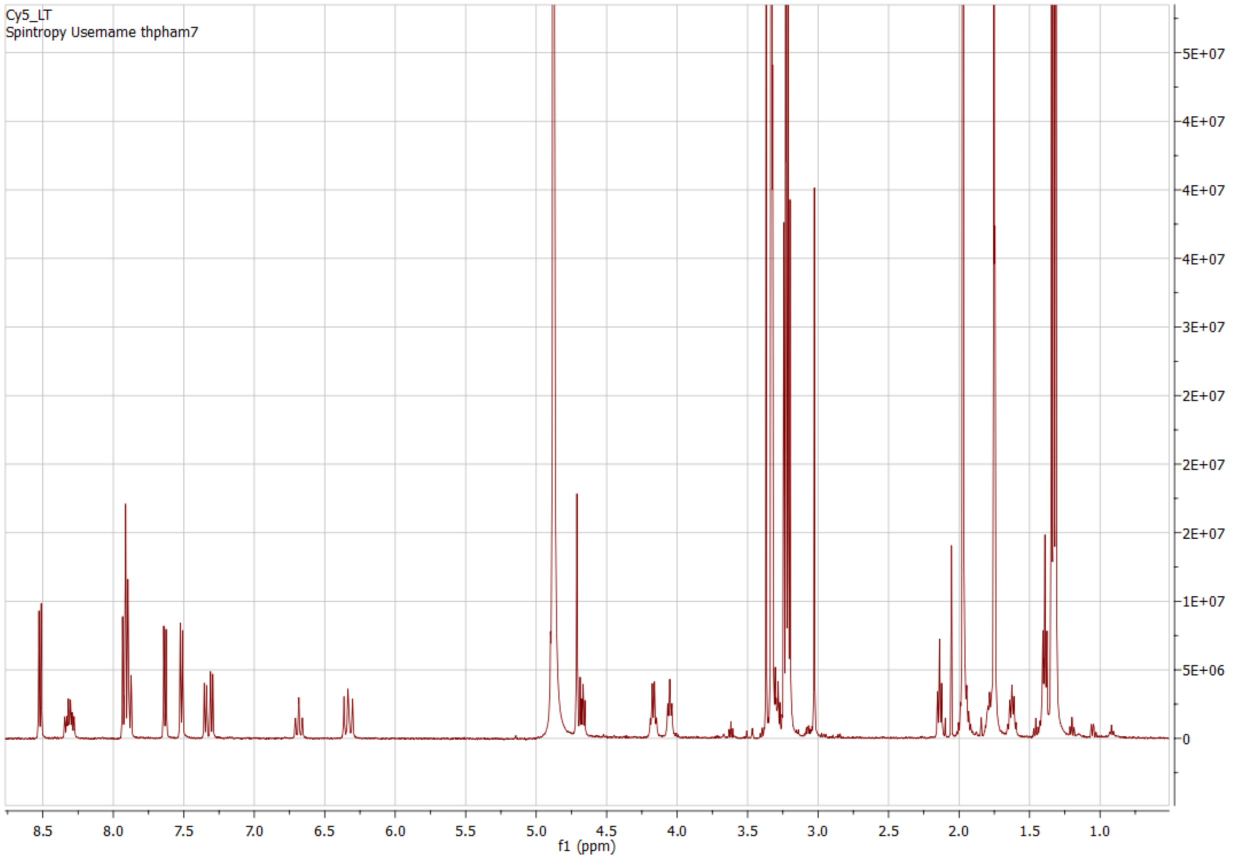


Figure S6. 500MHz [1]H NMR spectra of Tetrazine-$N_3$-Cy5 in $CD_3OD$

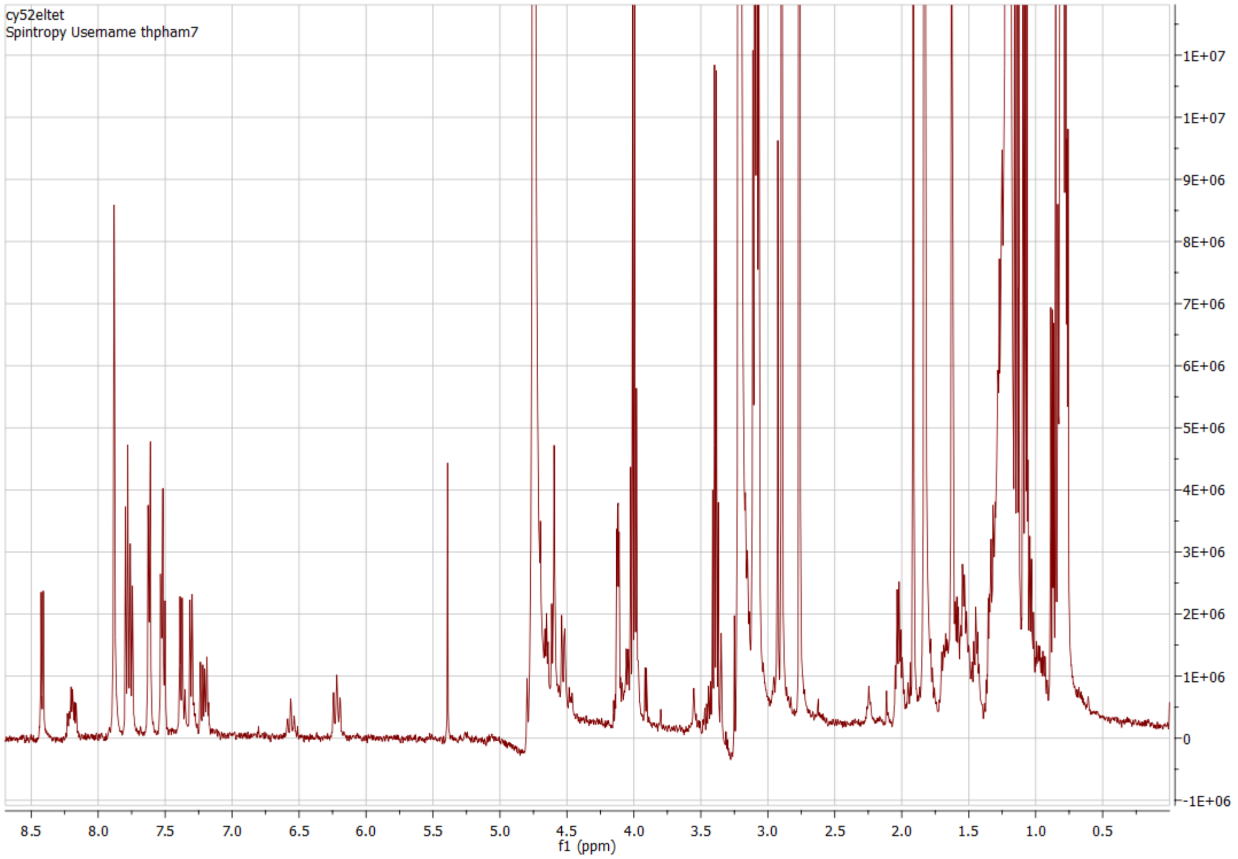


Figure S7. 500MHz $^1H$ NMR spectra of Tetrazine-$N_3$-$N_3$-Cy5 in $CD_3OD$

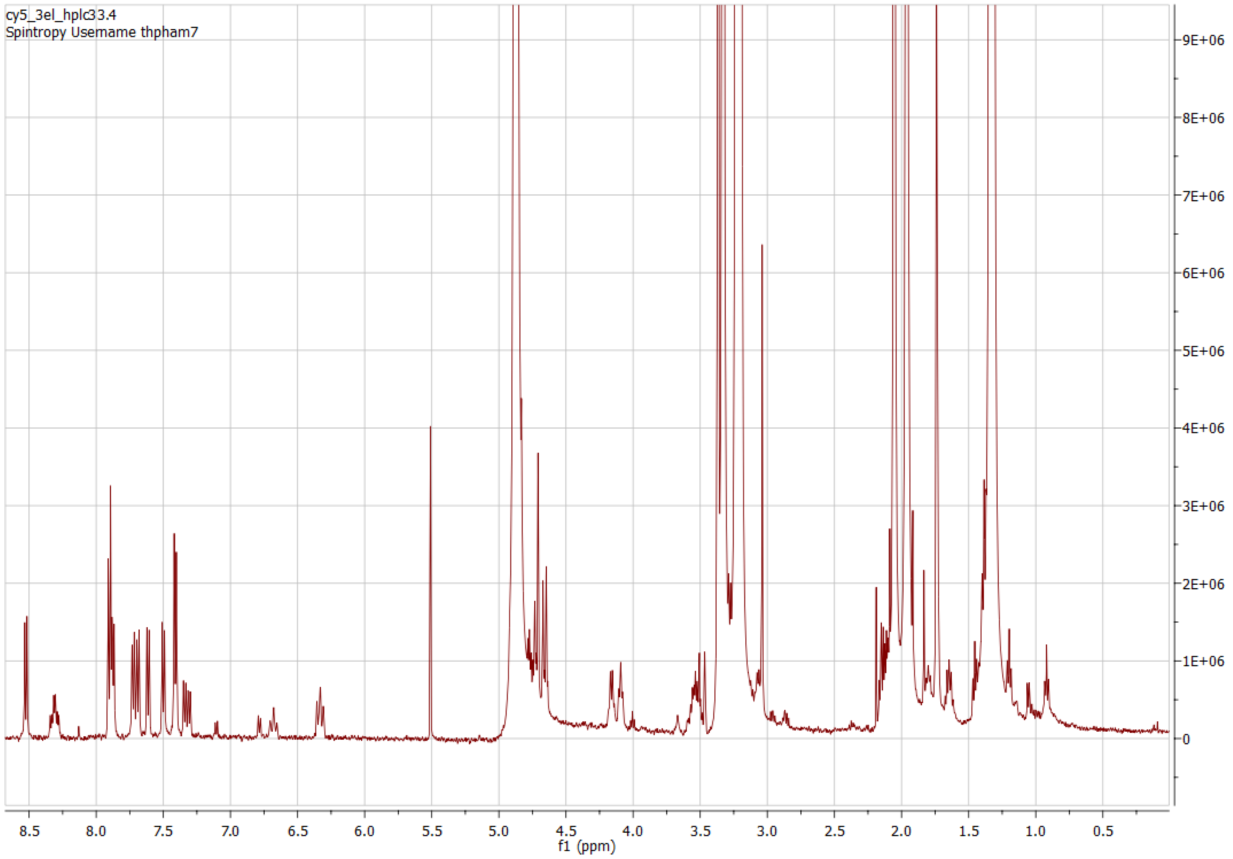


Figure S8. 500MHz $^{1}H$ NMR spectra of Tetrazine-$N_3$-$N_3$-$N_3$-Cy5 in $CD_3OD$

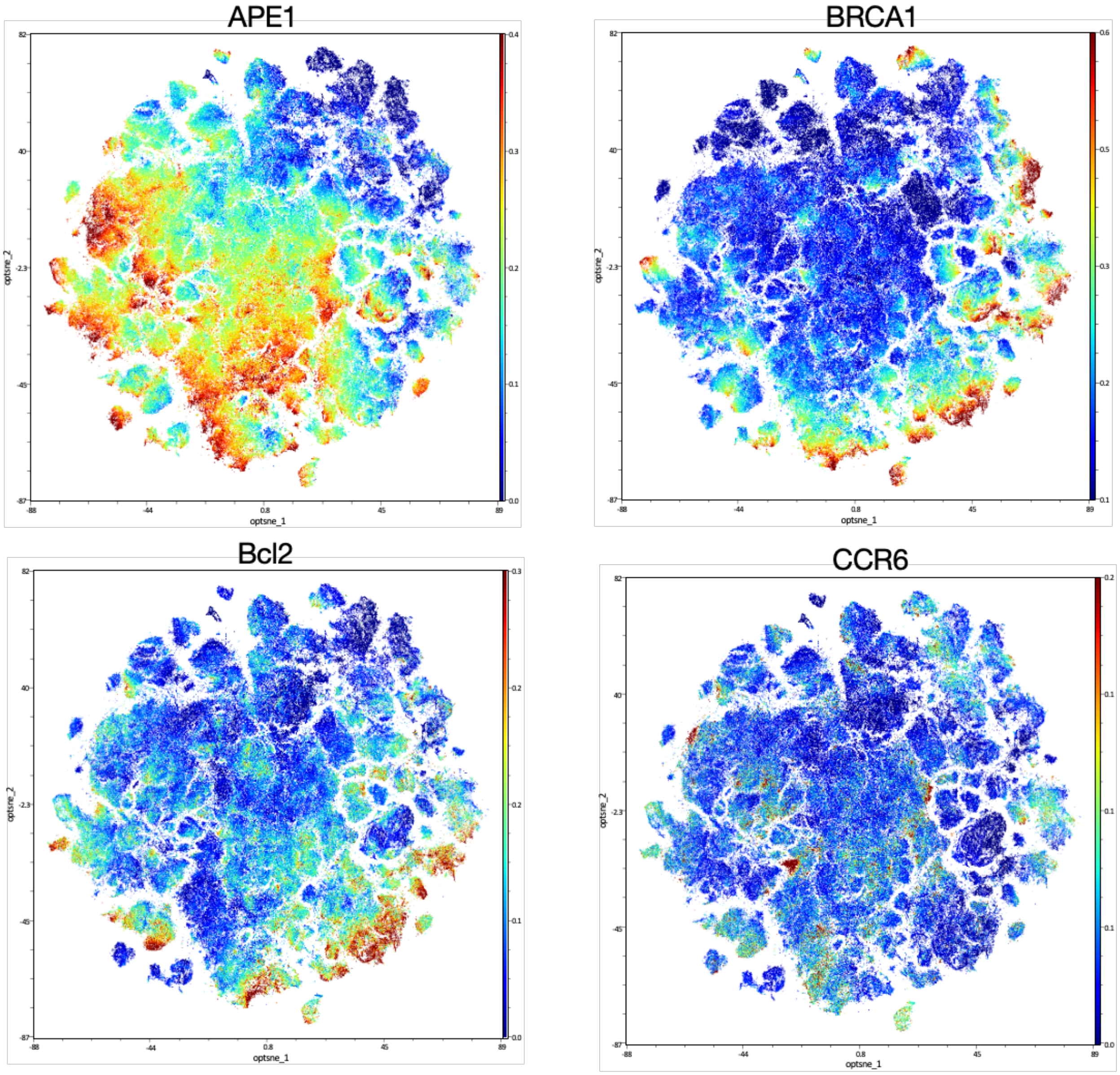
APE1
BRCA1
Bcl2
CCR6
optsne_1
optsne_2

CD11c

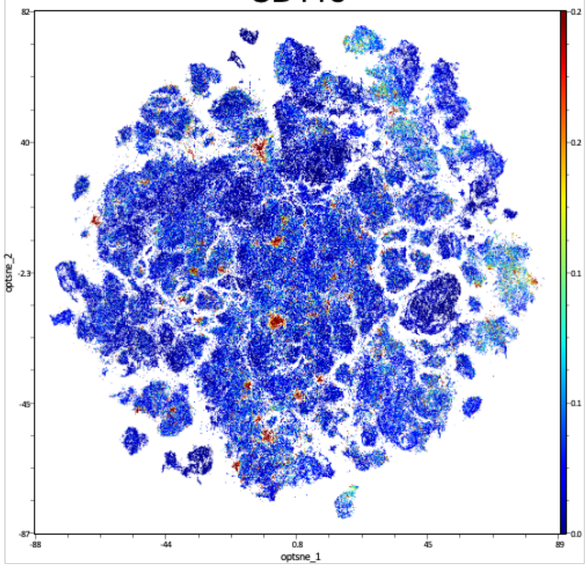


CD19

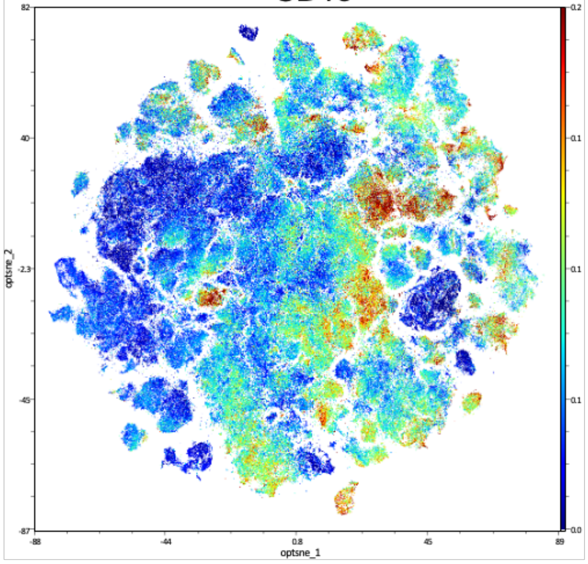


CD20

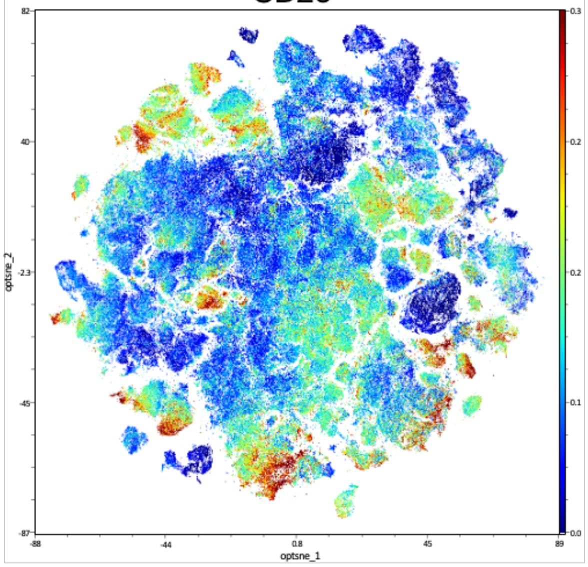


CD4

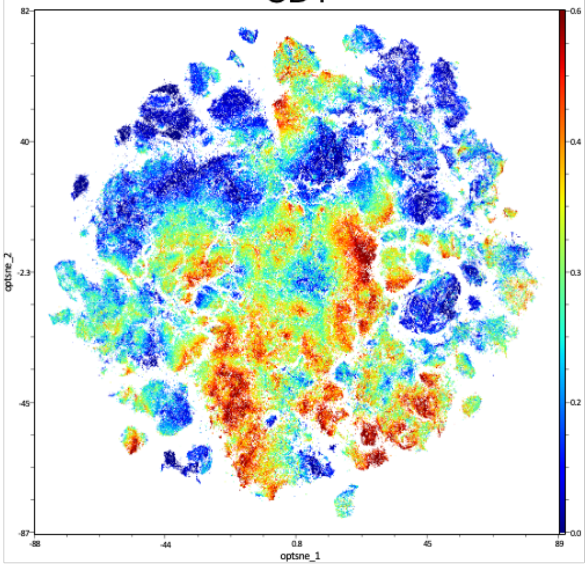

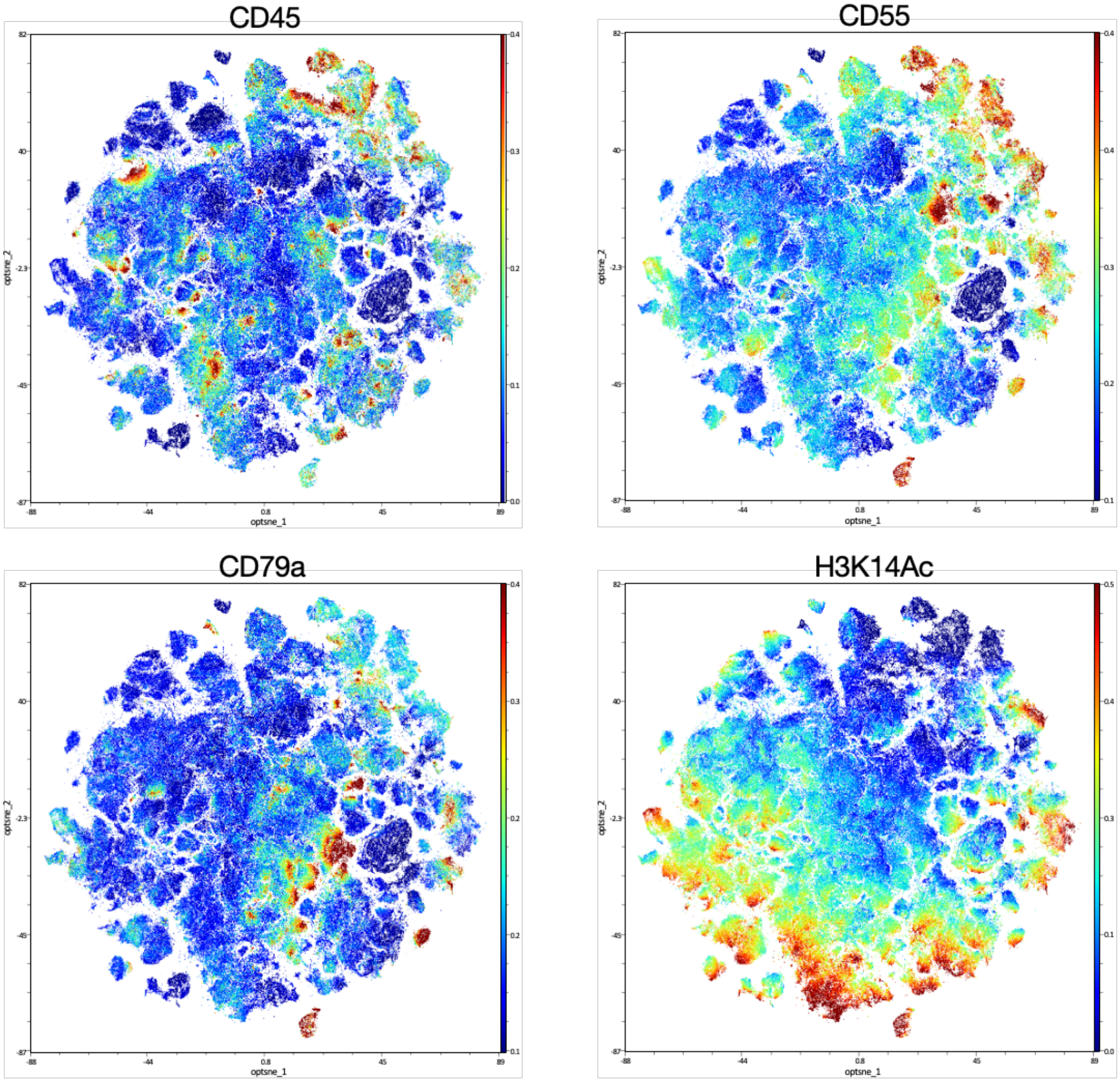
CD45
CD55
CD79a
H3K14Ac
optsne_1
optsne_2
82
40
-2.3
-45
-87
-88
-44
0.8
45
89

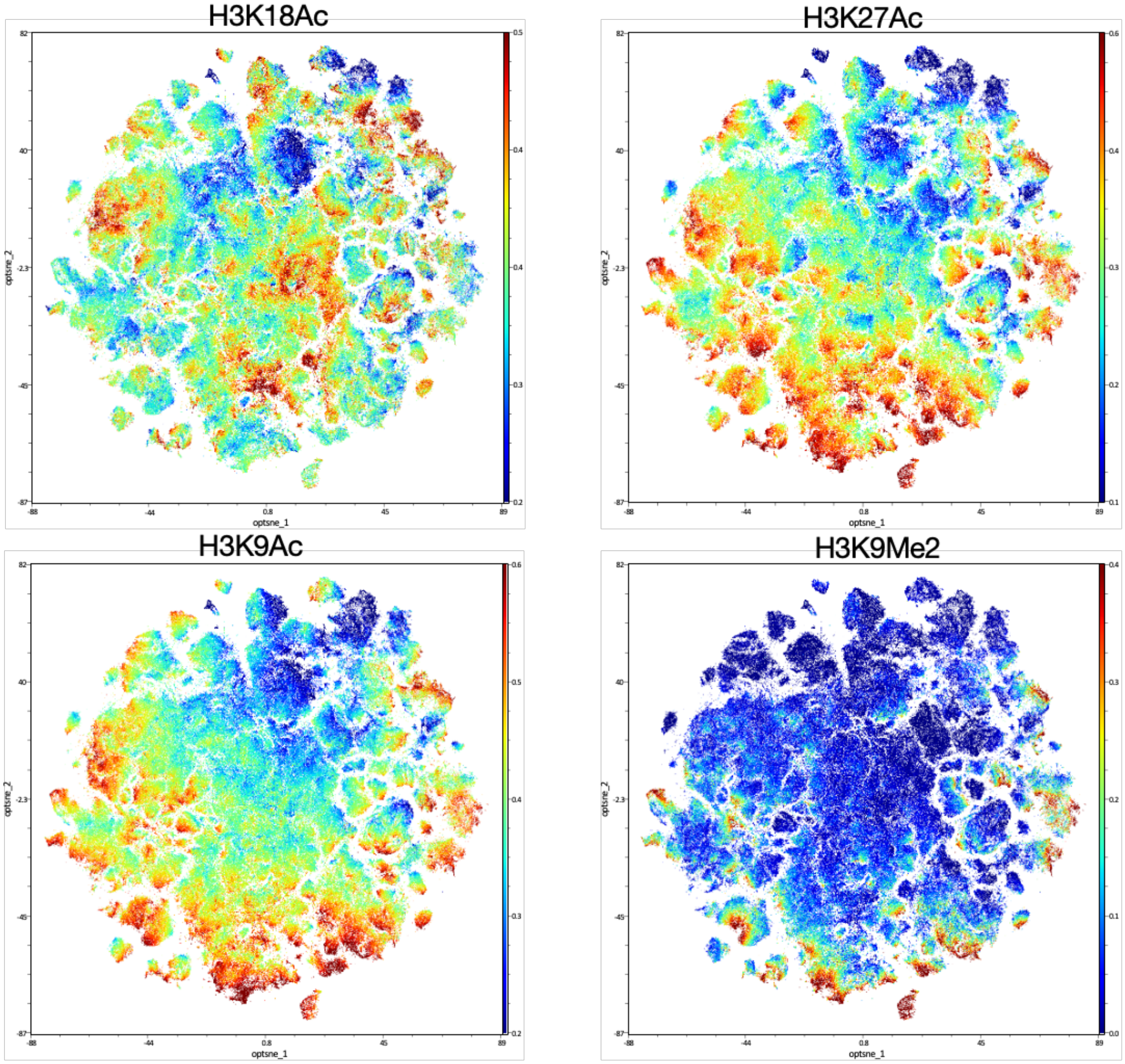
H3K18Ac
H3K27Ac
H3K9Ac
H3K9Me2
optsne_1
optsne_2

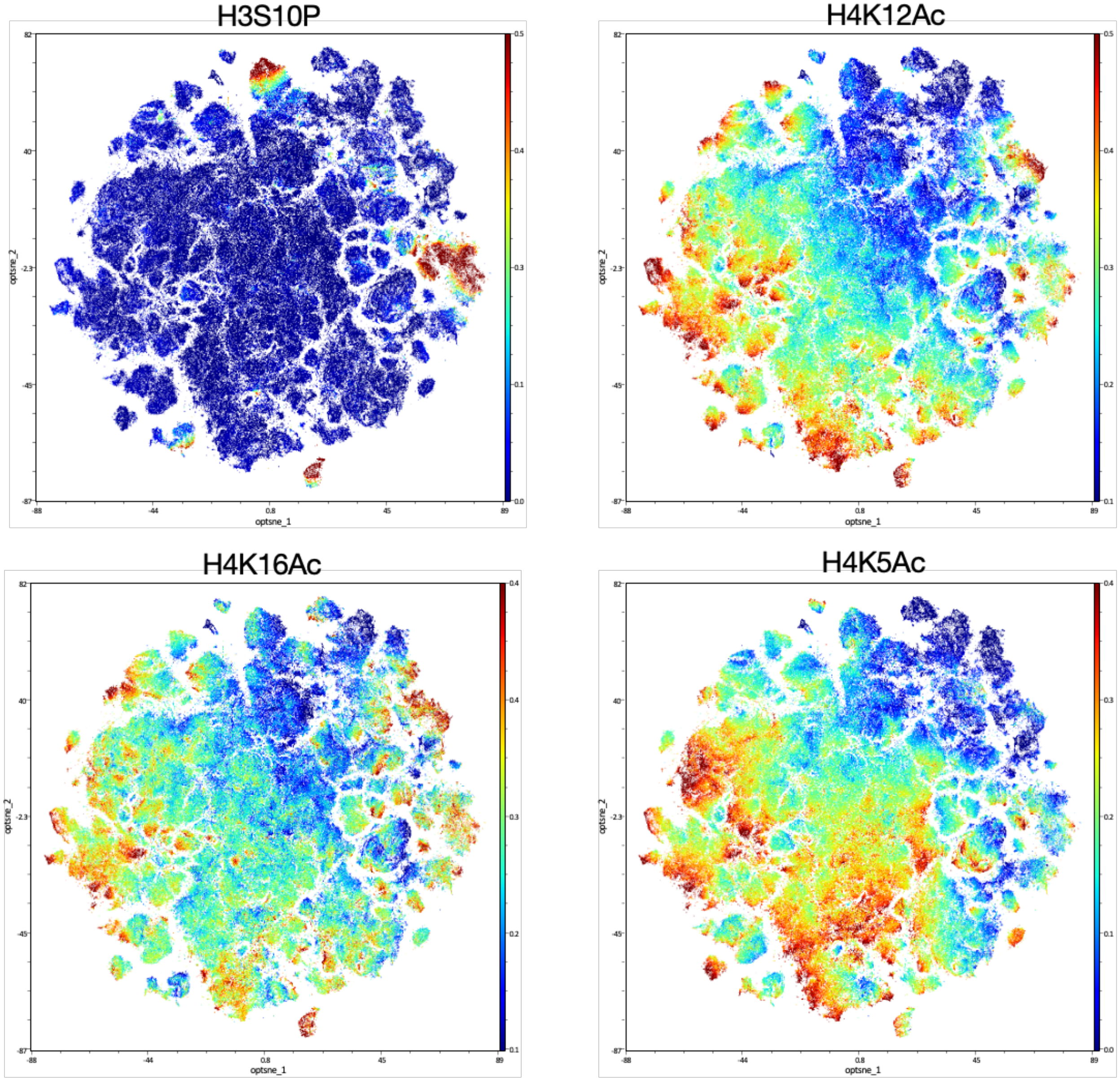

H3S10P
H4K12Ac
H4K16Ac
H4K5Ac
optsne_1
optsne_2

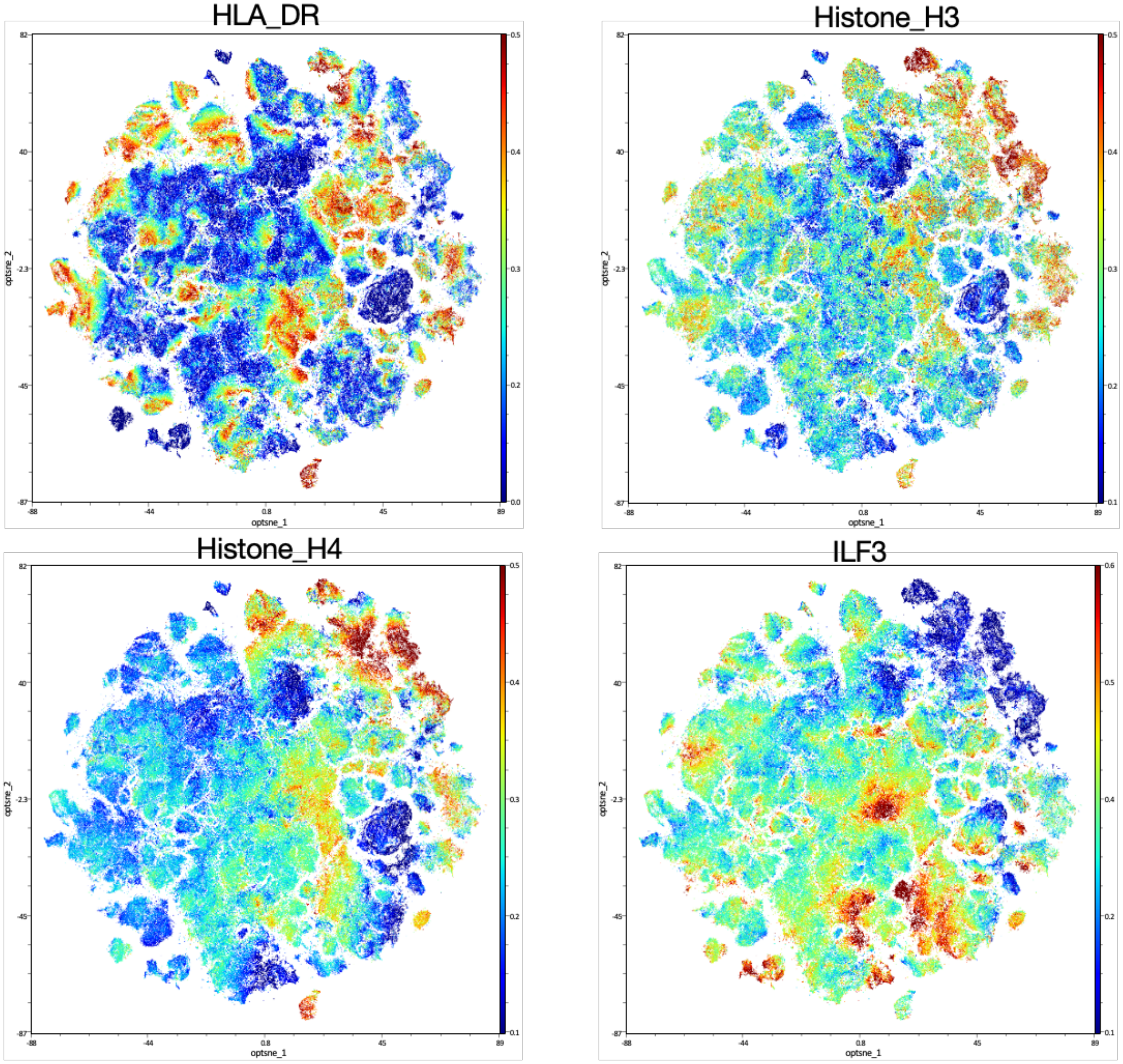
HLA_DR
Histone_H3
Histone_H4
ILF3
optsne_1
optsne_2

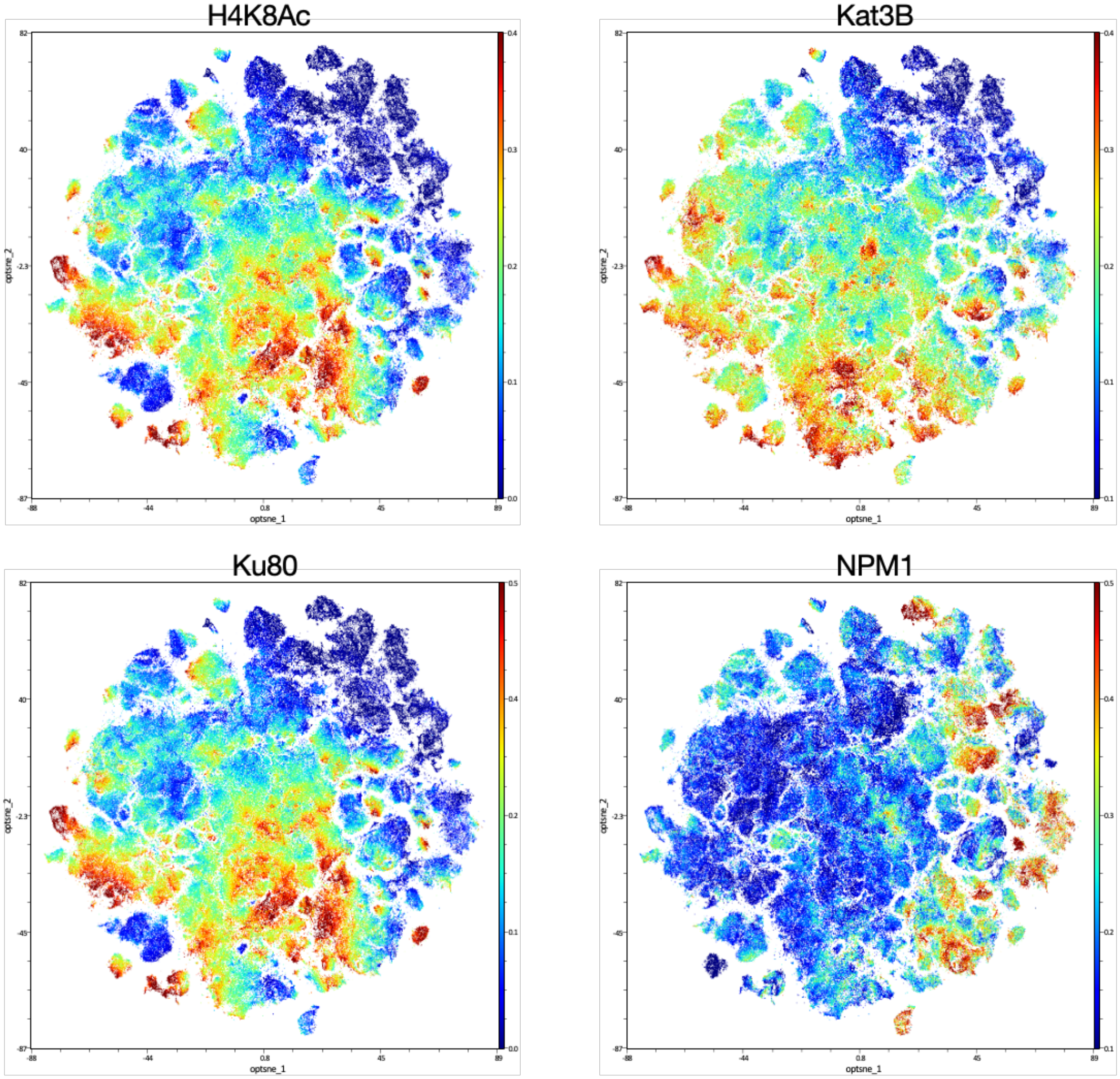


Figure S9. Single cell protein expression distribution in Optsne plots.

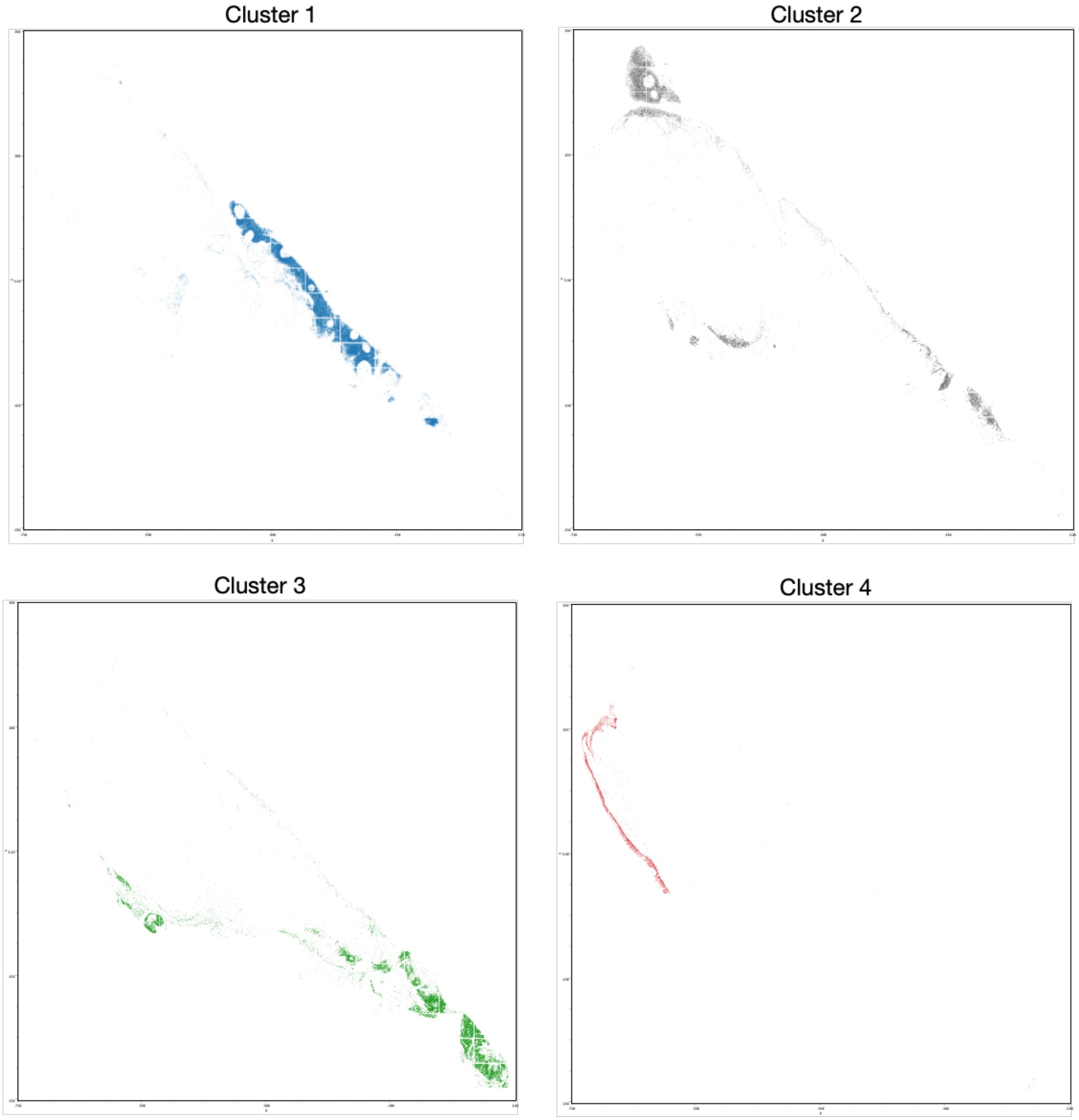
Cluster 1
Cluster 2
Cluster 3
Cluster 4

Cluster 5

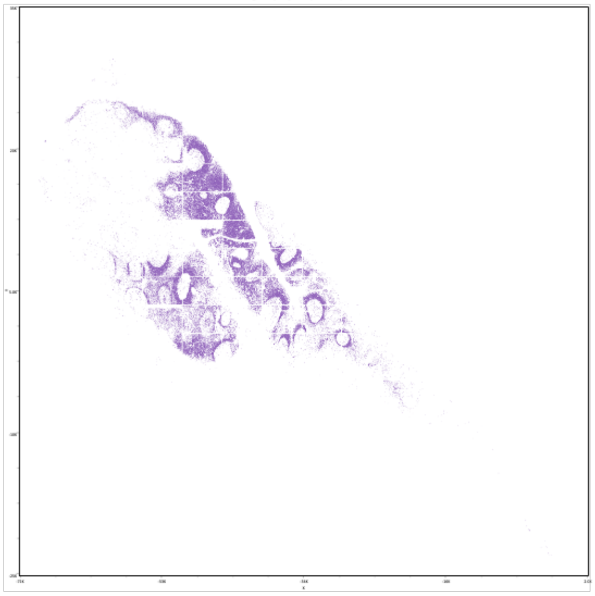

Cluster 6

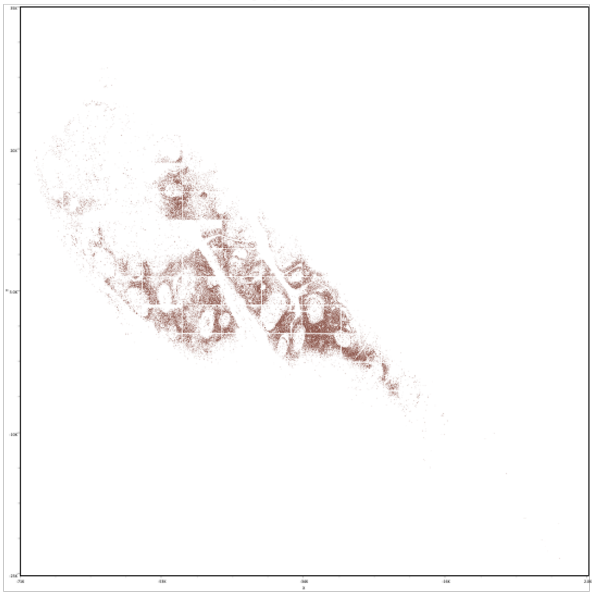

Cluster 7

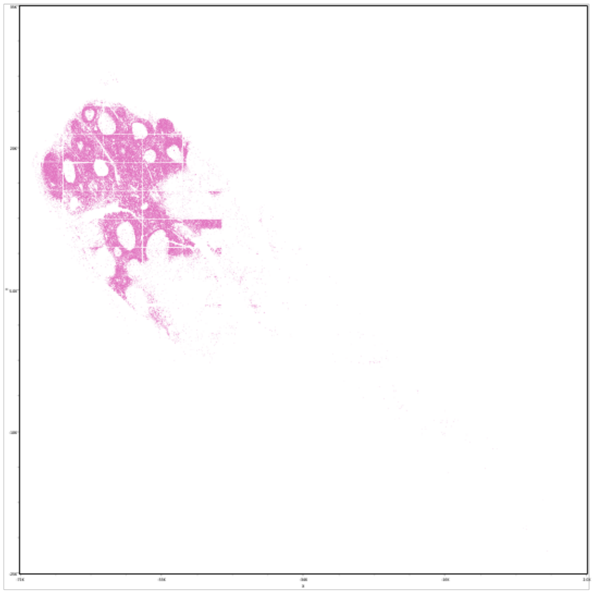

Cluster 8

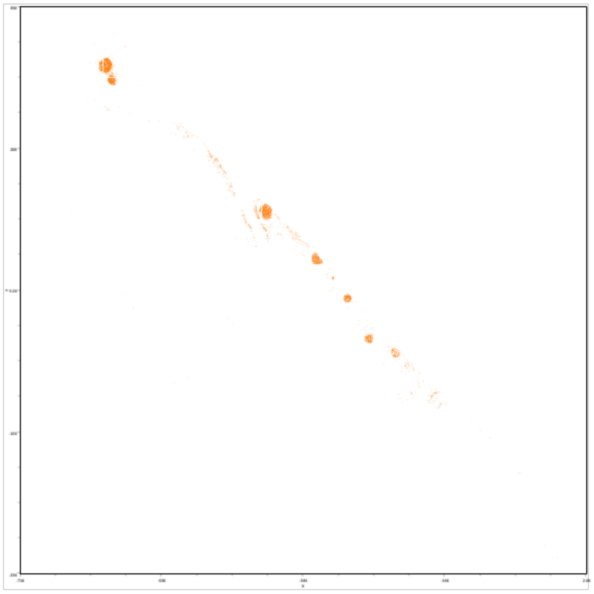

Cluster 9

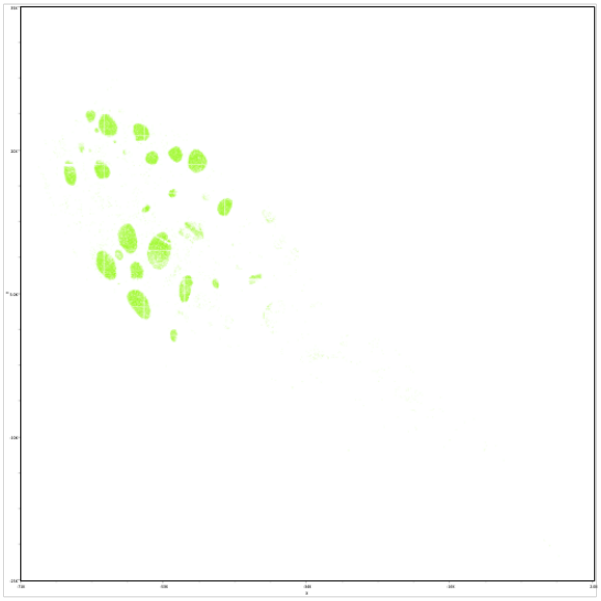

Cluster 10

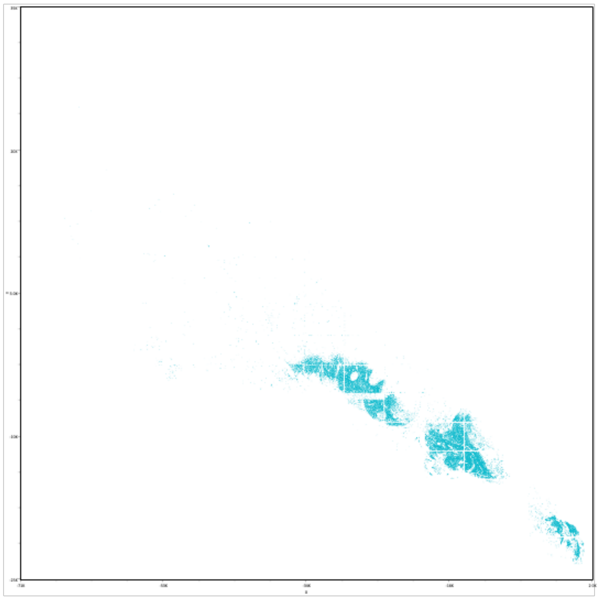

Cluster 11

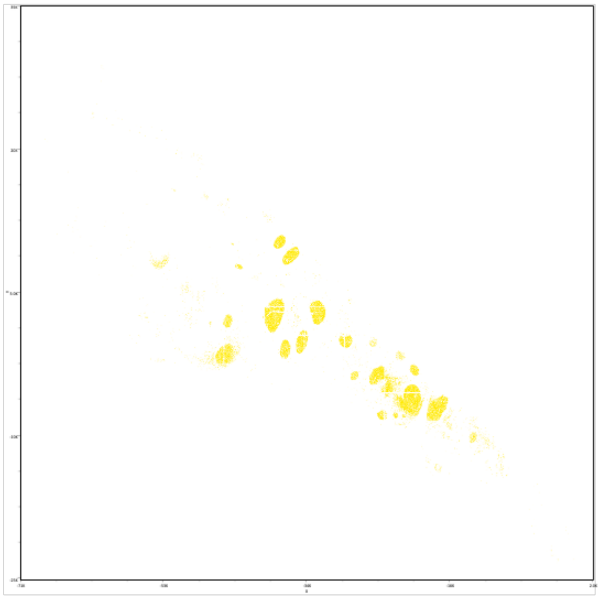

Cluster 12

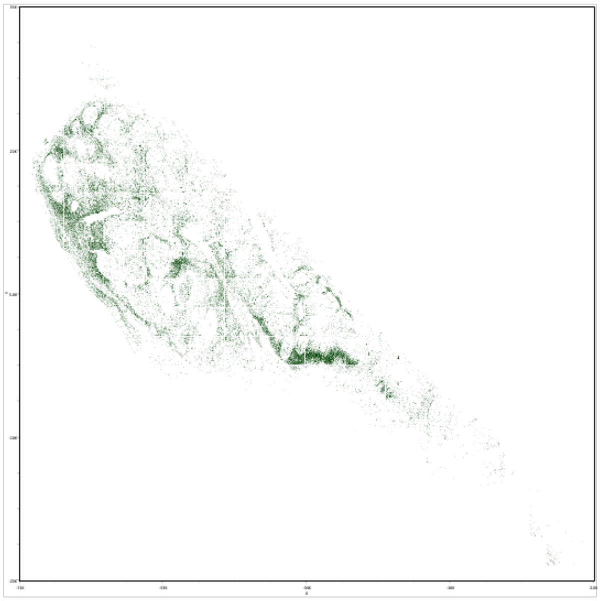

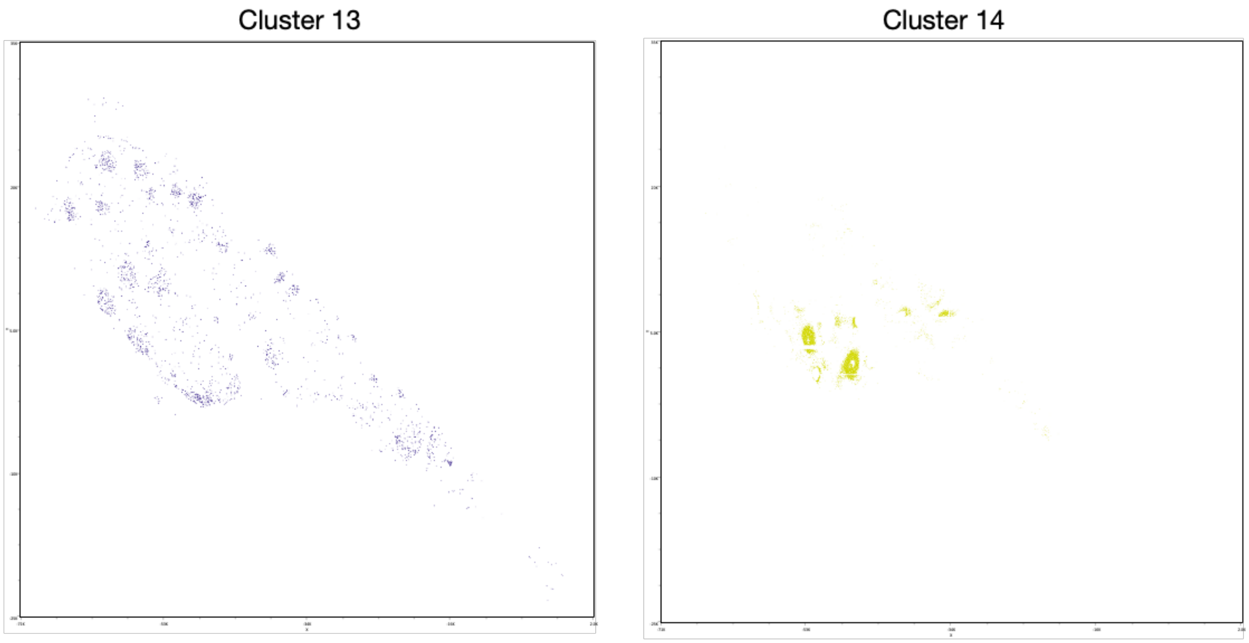


Figure S10. Anatomical locations of the individual cells from different cell clusters.
